# Data-driven mathematical modelling explains altered timing of *EARLY FLOWERING 3* in the wheat circadian oscillator

**DOI:** 10.1101/2025.04.08.644541

**Authors:** Abhishek Upadhyay, Jamila Rowland-Chandler, Julia Stewart-Wood, Gabriela Pingarron-Cardenas, Isao T Tokuda, Alex A R Webb, James C W Locke

## Abstract

Circadian rhythms, daily oscillations with a free-running period of approximately 24 hours, have evolved in organisms across the kingdoms of life, enabling organisms to anticipate and adapt to environmental cycles. Circadian timing in plants is governed by an oscillator gene network of transcriptional regulators that exists in each cell. Wheat provides an opportunity to investigate the mechanisms of the plant circadian oscillator in an important agricultural species. We recently found that a single oscillator component *EARLY FLOWERING 3* is expressed at a different time in wheat compared to the model plant Arabidopsis. This was unexpected since there is remarkable conservation of timing of activity of the different oscillator components within a kingdom, for example, even when animals switch from nocturnal to diurnal activity. We have examined how the change in timing of *ELF3* transcription between Arabidopsis and wheat has occurred and its implications for circadian oscillator function. We describe an optimised computational model of the wheat circadian oscillator that is informed by experimental data and the structure of the promoter elements driving oscillator gene expression. Our optimised computational models suggest that the dawn-expression of the key oscillator gene *ELF3* in wheat occurs due to repression of the *ELF3* promoter by TOC1. Our simulations predict that plant circadian oscillators are robust against changes in *ELF3* timing. Our work demonstrates that plant circadian oscillators can have a flexible architecture such that different oscillator structures can originate circadian rhythms.

## 1. Introduction

Many living organisms, including bacteria, cyanobacteria, fungi, plants and animals, exhibit 24-hour circadian rhythms (Konopka & Benzer 1971); (Hardin *et al*. 1990); (Merrow *et al*. 1999); (Roenneberg *et al*. 2003); (Nakajima *et al*. 2005); (Mittag *et al*. 2005); (Sartor *et al*. 2023). These biological oscillations allow organisms to anticipate rhythmic environmental signals such as light, temperature, and nutrient availability. Circadian oscillators regulate a wide variety of molecular and physiological processes in a coordinated and sequential manner (Dibner *et al*. 2010); (Reddy & O’Neill 2010); (Haydon *et al*. 2013); (Partch *et al*. 2014); (Piro *et al*. 2023). Therefore, understanding the fundamental mechanisms governing circadian oscillators might have impacts in the fields of agriculture and medicine (Steed *et al*. 2021); (Kramer *et al*. 2022).

An iteration between experiment and modelling has helped elucidate the structure of the circadian oscillator in the model plant *Arabidopsis thaliana* (Nakamichi *et al*. 2010; Pokhilko *et al*. 2012); (Millar *et al*. 2016); (De Caluwé *et al*. 2016); (Fogelmark & Troein 2014). The oscillator is composed of interlocking transcriptional– translational feedback loops that generate rhythmic gene expression over a 24-hour cycle. A simplified view of the network reveals a sequential expression of transcriptional repressors from morning to night. At dawn, *CIRCADIAN CLOCK ASSOCIATED 1* (*CCA1*) and *LATE ELONGATED HYPOCOTYL* (*LHY*) are expressed and repress the transcription of daytime *PSEUDO RESPONSE REGULATOR* (*PRR*) genes, including *PRR9, PRR7, PRR5*, and *PRR1* (also known as *TIMING OF CAB EXPRESSION 1, TOC1*) (Matsushika et al. 2000); (Nakamichi et al. 2010); (Huang et al. 2012); (Davis et al. 2022); (Wang et al. 2022). As the day progresses, the PRRs accumulate and feed back to repress *CCA1* and *LHY*, forming a core negative feedback loop.

As CCA1 and LHY protein levels diminish following their morning peak, the expression of *ELF3, ELF4*, and *LUX* increases, enabling assembly of the Evening Complex (EC) during the early night (Nusinow et al. 2011). This complex represses transcription of *TOC1* and other evening-phased genes. Concurrently, degradation of PRR proteins permits the reactivation of *CCA1* and *LHY* transcription later in the night, thereby resetting the oscillator for the next circadian cycle. These transcriptional dynamics are further regulated post-translationally via protein degradation mediated by factors such as the E3 ubiquitin ligase ZEITLUPE (ZTL) and GIGANTEA (GI). Together, these feedback loops generate a self-sustained oscillation with ∼24-hour periodicity. Orthologues of the key Arabidopsis oscillator components are present in major crops, where they have been shown to influence agriculturally important traits such as flowering time and yield (Steed et al. 2021).

The expression of the EC components in Arabidopsis are tightly coordinated, presumably to facilitate the formation of the complex, such that in diel light and dark cycles the transcripts of *ELF3, ELF4* and *LUX* all peak near dusk. In constant light, the timing of *ELF4* and *LUX* transcription remains locked to near subjective dusk, though *ELF3* transcript abundance peaks later through the night (Mockler *et al*. 2007). Recently, we reported that in wheat the peak timing of *ELF3* transcript abundance is at dawn, whereas *LUX* peaks at dusk as in Arabidopsis (Wittern *et al*. 2023). This anti-phased timing of the peak abundance of two of the components of the EC in wheat and other cereals raises question about the architecture of circadian oscillators in wheat, and the cereals in general (Alvarez *et al*. 2016); (Filichkin *et al*. 2011); (Lai *et al*. 2020); (Rees *et al*. 2022); (Yue *et al*. 2021). Circadian oscillator components regulate yield traits such as flowering and grain production (Wittern *et al*. 2023); (Taylor *et al*. 2024). Almost 1/5th of the world’s dietary nutrients are derived from wheat. Given the increasing global population, there will be a further need to increase wheat yield in the near future. Therefore, understanding the nature of circadian oscillators in wheat has practical implications.

Here we used a computational approach to model circadian oscillators in wheat, aiming to understand the timing differences in transcript accumulation among EC members. We adapted Arabidopsis circadian oscillator models using wheat experimental data and insight into the nature of the promoter elements regulating *ELF3* expression. We optimized two Arabidopsis circadian oscillator models, a minimalist model (De Caluwé et al. 2016) and another providing deeper mechanistic insight (Fogelmark & Troein 2014), to describe the relationships among the molecular components of the wheat circadian oscillator, with a specific focus on the timing, regulation, and potential functions of ELF3. Our analysis shows that the wheat circadian oscillator is likely to contain an additional link in which the evening-expressed TOC1 protein acts as a transcriptional repressor of *ELF*3. This repression can allow *ELF3* to be expressed in the morning, without shifting the phase of the other oscillator components. We observe that *ELF3* is dawn-phased under a range of photoperiods in our simulations, and test our model by comparing simulations of an *elf3* mutation to experimental data. Overall, we find that small perturbations to the circadian gene network of Arabidopsis can explain circadian dynamics in wheat.

## 2. Results

### 2.1. Arabidopsis circadian oscillator models do not capture wheat dynamics

Measurements of transcript abundance in long days (16 h light and 8 h dark) demonstrate that in Arabidopsis, *ELF3, ELF4* and *LUX* are co-expressed at dusk in LD cycles (Fig 1A). Meanwhile, in *Triticum turgidum* cv. Kronos, transcript abundance of the *ELF3* homolog, *TtELF3*, peaks at dawn, approximately 12 h after the peak in *TtLUX* transcript abundance near dusk (Fig 1B) .

**Figure 1.**
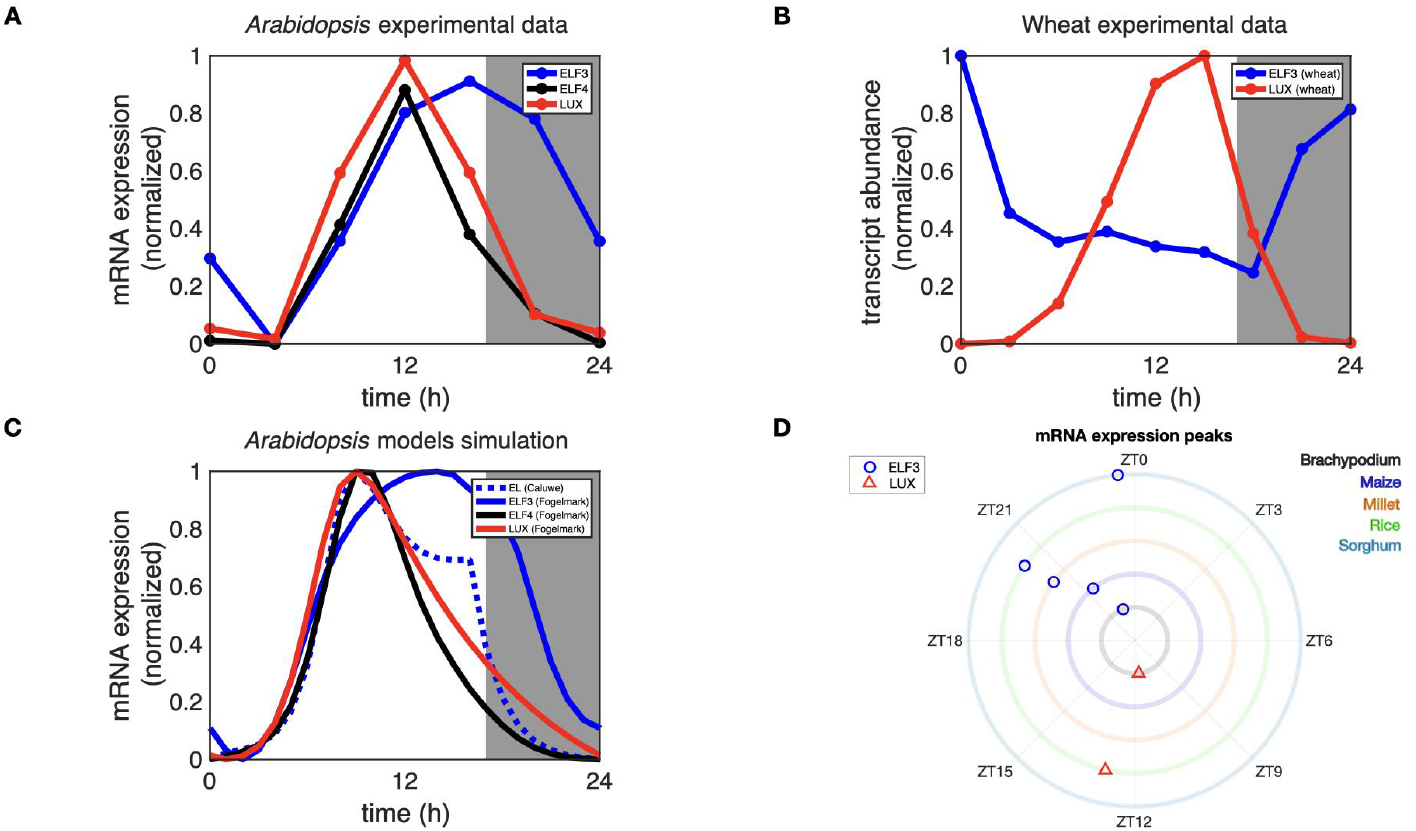
Current Arabidopsis circadian oscillator models do not capture dawn peaks of *ELF3* transcript abundance in cereals. **(A)** Measured dynamics of wild type Arabidopsis transcripts encoding EC proteins (adapted from (Mockler et al. 2007)). **(B)** Measured dynamics of *TtELF3* and *TtLUX* transcript abundance (from Wittern *et al*. 2023), White panels indicate day and grey panels indicate night. **(C)** Model simulations of Arabidopsis EC mRNA from de Caluwé 2016 and Fogelmark 2014. Blue, black and red solid lines are for *ELF3, ELF4* and *LUX*, respectively. Dashed blue line is the EC as represented in the reduced de Caluwe model. **(D)** Measured peak phases of *ELF3* (blue circle) and *LUX* (red triangle) among various cereals (denoted with circles of corresponding colours from inward to outward direction) are shown in polar plot. ZT stands for zeitgeber time and phases on a circle go clockwise from 0 to 24 hours.

Simulations of transcript abundance for EC components using two up-to-date Arabidopsis circadian oscillator models, the ‘compact’ model of de Caluwé 2016 and the more complete Fogelmark 2014 model (Fig 1C) predicted the measured Arabidopsis data well (Fig 1C), showing that these models can reproduce the dynamics of the evening complex in Arabidopsis. Unsurprisingly, these models did not capture the very different dynamics of the wheat EC components. We have not included *ELF4* in the analysis of wheat transcripts because the identity of a *bona fide* wheat *ELF4* that is a component of circadian oscillators is unclear (Wittern *et al*. 2023).

Morning phased expression of *ELF3* has been observed in other species of wheat, although one study in bread wheat (*Triticum aestivum*) reported an evening phased damped *ELF3* rhythm under constant conditions (Rees *et al*. 2022). To compare how evening complex genes peak among other cereals, we collected their phases from the published literature (Alvarez *et al*. 2016); (Filichkin *et al*. 2011); (Lai *et al*. 2020); (Rees *et al*. 2022); (Yue *et al*. 2021) (Fig 1D). We find that the dawn-phased expression of *ELF3* orthologues, anti-phased to a dusk-phased expression of *LUX* orthologues, is a common feature of the cereals. *ELF3* orthologues transcripts peak near dawn in sorghum, maize (Lai *et al*. 2020), *Brachypodium distachyon* and rice (*Oryza sativa*) (Filichkin *et al*. 2011), whilst *LUX* orthologues have peak transcript abundance near dusk (Fig 1D).

These data demonstrate that there is a fundamental difference in the patterning of circadian oscillators between the model Arabidopsis and the cereals. *ELF3* is required for circadian rhythms in wheat (Wittern *et al*. 2023) but current models of plant circadian oscillators do not describe its timing and regulation in wheat.

### 2.2 Two models of the wheat circadian oscillator

Our analysis of the regions upstream of the coding sequence of the wheat orthologs of *AtELF3* indicated the potential for TOC1-mediated regulation of *ELF3* expression, distinct from what we observed in the *A. thaliana* promoter (Wittern *et al*. 2023). In *A. thaliana, ELF3* expression is repressed by CCA1, which binds a CCA1-binding site (CBS) upstream of the 5’ UTR (Lu *et al*. 2012) (Figure 2C, see Fig S1). Given that CCA1 and LHY also bind the *ELF3* promoter at the evening element (EE, AAAATATCT, (Harmer *et al*. 2000); (Alabadí *et al*. 2001); (Harmer & Kay 2005) in Arabidopsis and a short evening element (SEE, AAATATCT, Gong et al., 2022) in *T. aestivum*, we also searched the Arabidopsis and *T. aestivum* sequences for these motifs and found that the sequence upstream of *T. aestivum ELF3-D1* contains one SEE (Figure 2C). However, the *T. aestivum* genomic region upstream of the 5’ UTR does not contain the CBS, the motif in the Arabidopsis *ELF3* promoter bound by CCA1, in any of the A, B, or D homeologs. Instead, the sequences upstream of these wheat *ELF3* orthologs contain multiple motifs associated with TOC1 recruitment in Arabidopsis. These motifs contain a T1ME motif (TGTG, (Gendron *et al*. 2012) and include the morning element (ME, GTGTGG, (Gendron *et al*. 2012); (Harmer & Kay 2005), hormone-upregulated-at-dawn element (HUD, CATGTG, (Michael *et al*. 2008), and CO-response element (COR, TGTG(N2-N3)ATG, (Tiwari *et al*. 2010); (Gendron *et al*. 2012) (Figure 2C). Residues essential for binding the T1ME motif in AtTOC1 (ARGF within the CCT domain) are conserved in its wheat ortholog, indicating that TOC1 in wheat can likely bind T1ME-containing motifs (Gendron *et al*. 2012). We found similar TOC1 binding sites in the promoter of *ELF3* orthologs in other cereals (Fig S1).

**Figure 2.**
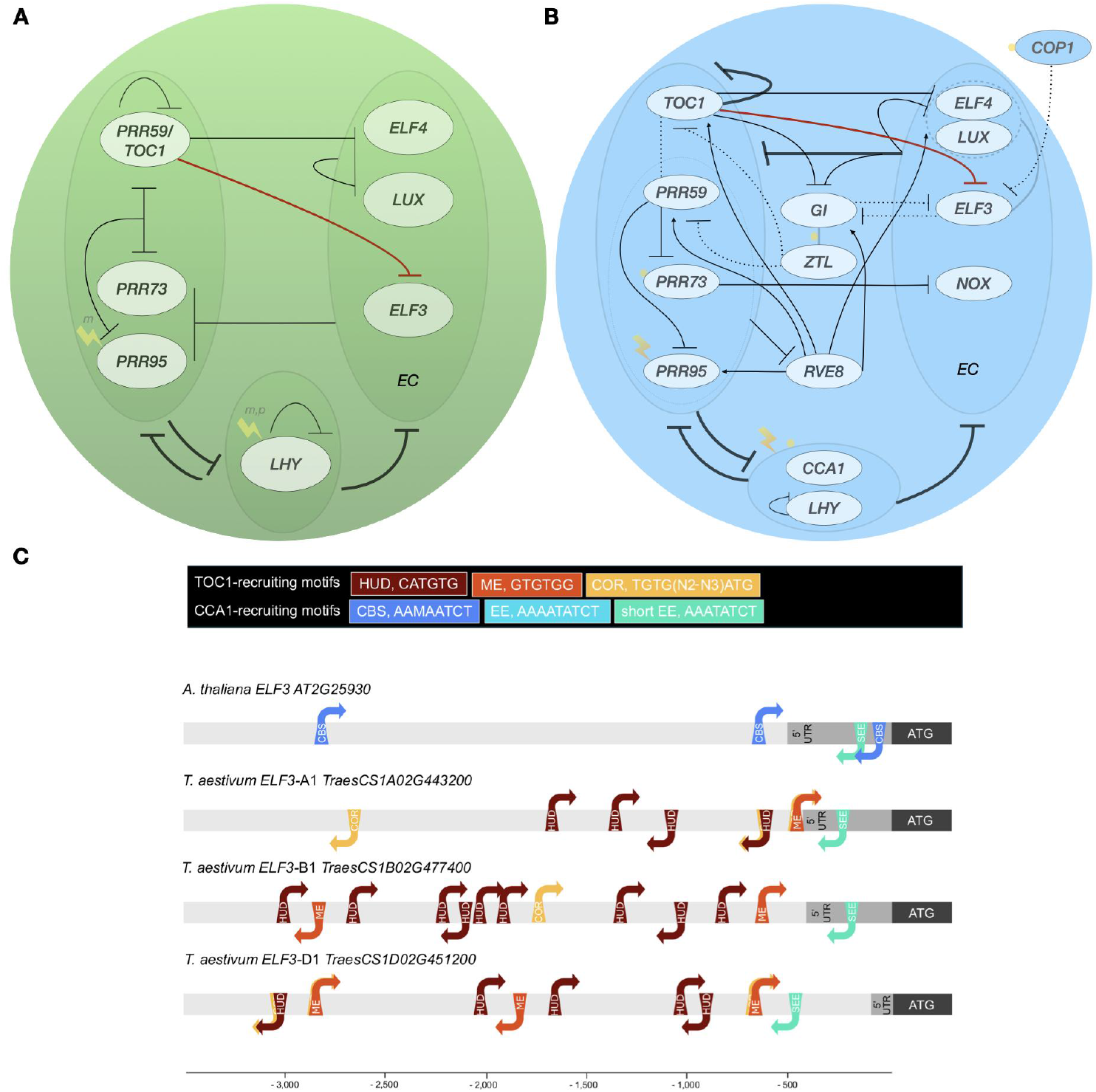
Schematic representations of the compact (A) and large (B) mathematical models of the wheat circadian oscillator. The network diagrams illustrate the oscillator network, which includes dawn (*LHY* or *LHY/CCA1*), daytime (*PRRs/TOC1*), and evening/night (the EC) classes of genes shown in large ovals with lighter boundaries. Specific genes are shown in small ovals with dark boundaries. Light input at transcriptional level is represented with a yellow lightning arrow and at translational level by a yellow filled circle. In the compact model protein degradation rates can also be altered between dark and light (not depicted, see Supplementary section 1). In the large model, solid lines represent transcriptional regulation and dashed lines depict protein– protein interactions. Red repression curve line in both models denote our assumption of repression of *ELF3* by TOC1. **(C) *AtELF3* and *TaELF3* promoter region analysis**. Representation of genomic DNA sequences 3500 base pairs upstream of the start codon of *AtELF3* and its orthologs in *T. aestivum*. Motifs associated with TOC1 recruitment include the morning element (ME, GTGTGG), hormone-upregulated-at-dawn element (HUD, CATGTG), and CO-response element (COR, TGTG(N2-N3)ATG). Motifs associated with CCA1 recruitment include the CCA1 binding site (AAAAATCT or AACAATCT), evening element (EE, AAAATATCT) in Arabidopsis and shortened evening element (SEE, AAATATCT) in wheat. Notably, the promoter sequences of wheat *ELF3* orthologs include elements associated with TOC1 binding that are not present in the promoter region of the *A. thaliana* gene. We hypothesize that TOC1 represses expression of *ELF3* in wheat but not in *A. thaliana*, resulting in morning-phased expression of *ELF3*.

While regulation of *ELF3* by TOC1 has not been reported in Arabidopsis, the abundance of binding sites for TOC1 in the promoters of wheat *ELF3* orthologues (Figure 2C) led us to test the hypothesis that the altered timing of *ELF3* expression in wheat compared to *AtELF3* is a consequence of TOC1 repression of *ELF3* in wheat. This repression is not described in models of Arabidopsis circadian oscillators. We therefore developed two models of circadian oscillators in wheat by incorporating a presumed regulation of *ELF3* transcription by TOC1 into extant models of *A. thaliana* circadian oscillators (Fogelmark & Troein 2014); (Greenwood *et al*. 2022).

The first model (see Fig 2A) we chose for development is a compact model consisting of 9 equations and 34 parameters where oscillator components are combined to provide a minimal description of the oscillatory network (De Caluwé *et al*. 2016); (Greenwood *et al*. 2022). The second model (see Fig 2B) is a detailed representation of biochemical interactions within the circadian oscillator, optimised using a large dataset. It comprises 35 variables and 119 parameters (Fogelmark & Troein 2014). Details of equations, parameters, and model development for the wheat models are described in the methods section 4.2.

### 2.3 Model optimization explains the experimentally observed behaviour of wheat *ELF3*

Next, we optimised the parameters of both the compact and large wheat models to fit to experimental data on wheat oscillator gene transcript abundance. We fitted our models to our recently published dataset for six core wheat clock genes, including *TtELF3* and *TtLUX* (Supplementary Section 3, Fig S2 and Fig S3; (Wittern *et al*. 2023)). We tested whether we could capture wheat expression dynamics by only optimising the parameters related to *TtELF3* in the model. The details of the optimised parameters are described in Tables S1 and S2.

We fitted our model to the peaks and troughs of the experimental dataset using a Multiple-Objective Genetic Algorithm (Supplementary Section 3). The simulated dynamics using the optimal parameter set (Tables S1 and S2) closely aligned with the experimental data, with peak times within 1.30 and 2.13 hours of the data for compact and large models, respectively (Fig 3). Notably, the model now captures the dawn-expressed dynamics of *TtELF3* as well as the dusk-peaking behaviour of *TtLUX* (Figs 3 and 4).

**Figure 3.**
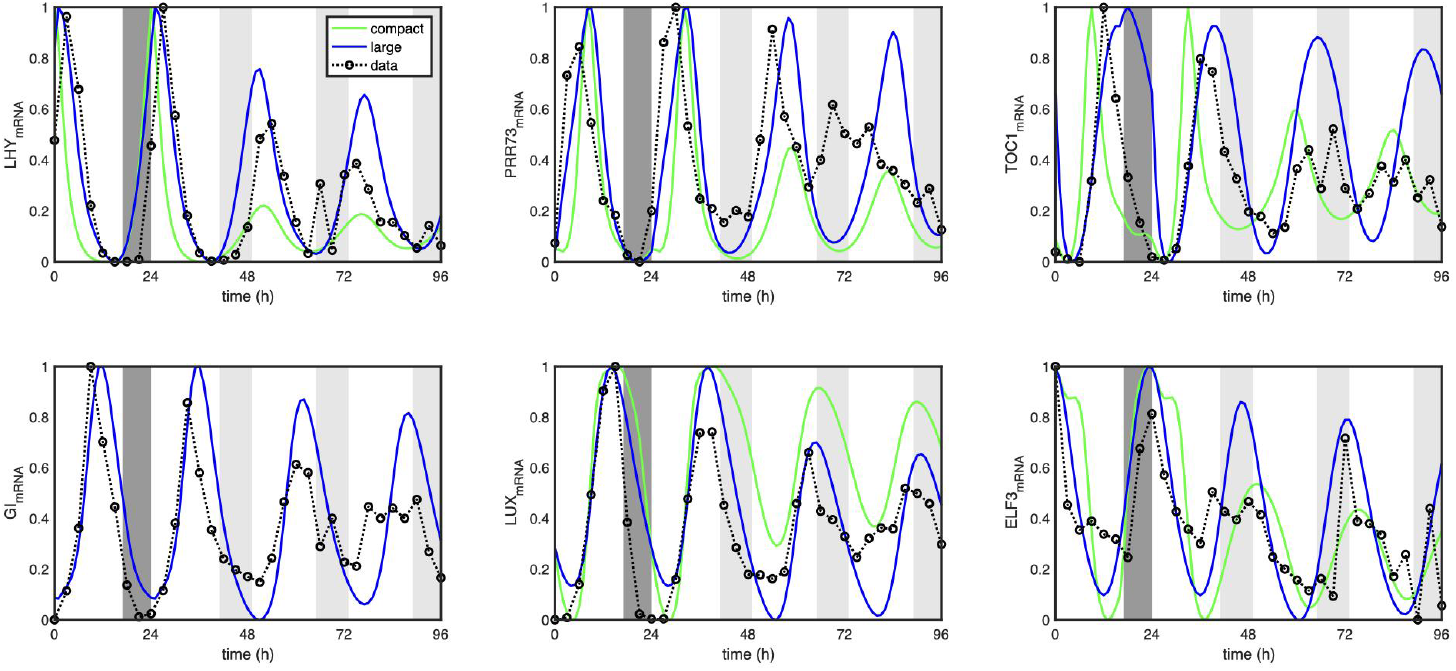
Optimised wheat models capture transcript abundance dynamics of key circadian oscillator genes. Post-optimization simulated dynamics of circadian oscillator genes are shown for both the compact (green solid lines) and large (blue solid lines) models, along with experimental data (black dotted line). The white panels represent daylight, while the dark grey panels indicate night during the first 24 hours. From 24 to 96 hours under constant light conditions, the white panels represent subjective day, and the lighter grey panels indicate subjective night. Both simulations and experimental data are normalised for all six genes.

**Figure 4.**
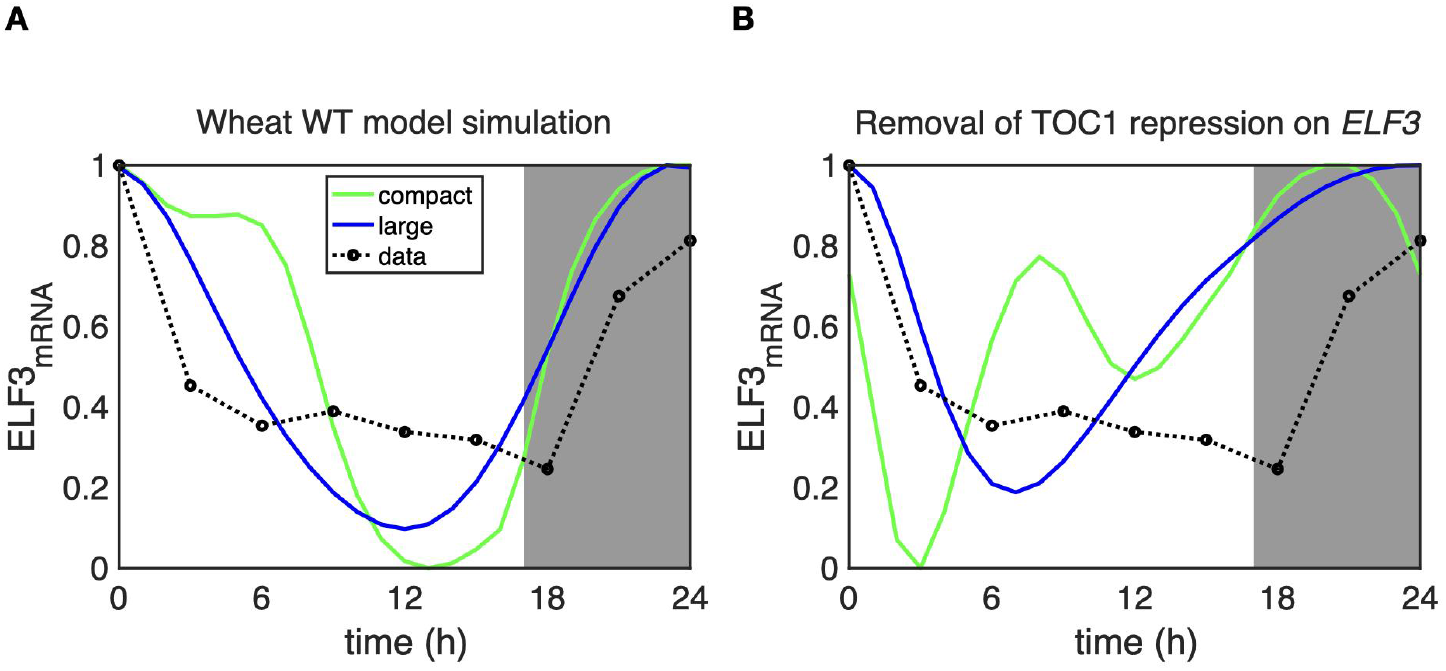
TOC1 repression of *ELF3* is required for correct timing of *ELF3* expression. **(A)** The experimental dynamics of wild type wheat *TtELF3* are replicated in simulations when TOC1-mediated repression of Tt*ELF3* is included. Simulations are depicted for compact (green solid lines) and large (blue solid line) models and corresponding experimental wild type data are plotted under long days (16:8) condition (black dotted line). The white panel mimics the light in day for 16 hrs and dark in grey panels for 8 hrs in night over the first 24 hr. The trough timings of wild type *ELF3* in the compact model is 13 hr and large is 12 hrs. **(B)** Simulations of *TtELF3* mRNA no longer match experiment when TOC1 repression of *TtELF3* is removed. When TOC1 repression is not applied, the trough timings of *TtELF3* in the compact model is 3 hr and in the large is 7 hrs.

We next examined ELF3 protein dynamics, as a recent paper has proposed that ELF3 protein is degraded in the light (Fig S4, (Alvarez *et al*. 2023)). Reproducing these findings, in the large model simulations of ELF3 protein levels peak at dawn in light dark cycles due to COP1-mediated degradation of ELF3. Further, if we increased the strength of the COP1-mediated degradation, we could observe even sharper drops of ELF3 protein in light without significantly affecting the phase of the other clock components (Alvarez *et al*. 2023) (Fig S4). This suggests that ELF3 degradation in light can be understood via our large model without the need for further components. In the wheat compact model, we did not include light mediated degradation of ELF3 so we did not observe a sharp drop in ELF3 protein at dawn (Fig S4). However, when we incorporated light mediated degradation we did observe ELF3 to drop faster at dawn in the compact model (Fig S4), suggesting this can be added to future models.

### 2.4 TOC1 repression of *ELF3* is required to explain the dynamics of wheat circadian oscillators

We tested whether our model addition of TOC1 repressing *ELF3* was required for dawn peaks of *ELF3* transcript abundance. If we removed the *ELF3* repression by TOC1, the dawn peak was no longer apparent (Fig 4). Furthermore, we were unable to recapture the dawn peak of *ELF3* when we re-optimised the model parameters without including TOC1 repression of *ELF3* (Fig S5). Conversely, if we removed *ELF3* repression by CCA1 in the model our simulations were largely unaffected (Fig S6), fitting with the lack of CCA1 binding sites in the *TtELF3* promoter (Fig 2C). This suggests that our hypothesis, based on the analysis of genomic sequences upstream of the ELF3 coding region, is correct: the wheat circadian oscillator functions differently compared to the Arabidopsis model due to TOC1 repressing *ELF3*.

### 2.5 Simulated *ELF3* expression tracks dawn over a range of photoperiods

We next tested whether *ELF3* peaks at dawn across a range of simulated photoperiods (i.e. duration of light during a day), or if this was specific to entrainment under long-day conditions (Figure 3). The photoperiod is a key environmental cue that can affect the phase of the circadian oscillator and its outputs, as well as the timing of many plant developmental processes such as flowering. We examined the phase timing of the EC components (ELF3, ELF4 and LUX) in simulations of the compact and large wheat and Arabidopsis models under seven different photoperiods. The simulated photoperiods were 3 h (3 h light and 21 h darkness), 6 h, 9 h, 12 h, 15 h, 18 h and 21 h (21 h light and 3 h darkness). *TtELF3* peaks at dawn in the compact wheat model (blue circles, Fig 5A) and near dawn in the large wheat model (blue circles, Fig 5B) for all photoperiods simulated. Moreover, *TtELF4* and *TtLUX* (black squares and red triangles, respectively) peak around the evening under all photoperiods in both wheat models. See Fig S7 for example wheat oscillator waveforms under 3 photoperiods. Our results contrast with the simulations of the Arabidopsis models. In the Arabidopsis simulations, all of the evening complex genes peak around dusk, although we note in the large Arabidopsis model (Fogelmark & Troein 2014) under long photoperiods (18 h-21 h light) the evening complex genes peak earlier in the day. These results suggest that *ELF3* peaks earlier in wheat than Arabidopsis across all photoperiods. This is of particular interest given that the timing of *ELF3* expression affects flowering time in wheat (Peng *et al*. 2024).

**Figure 5.**
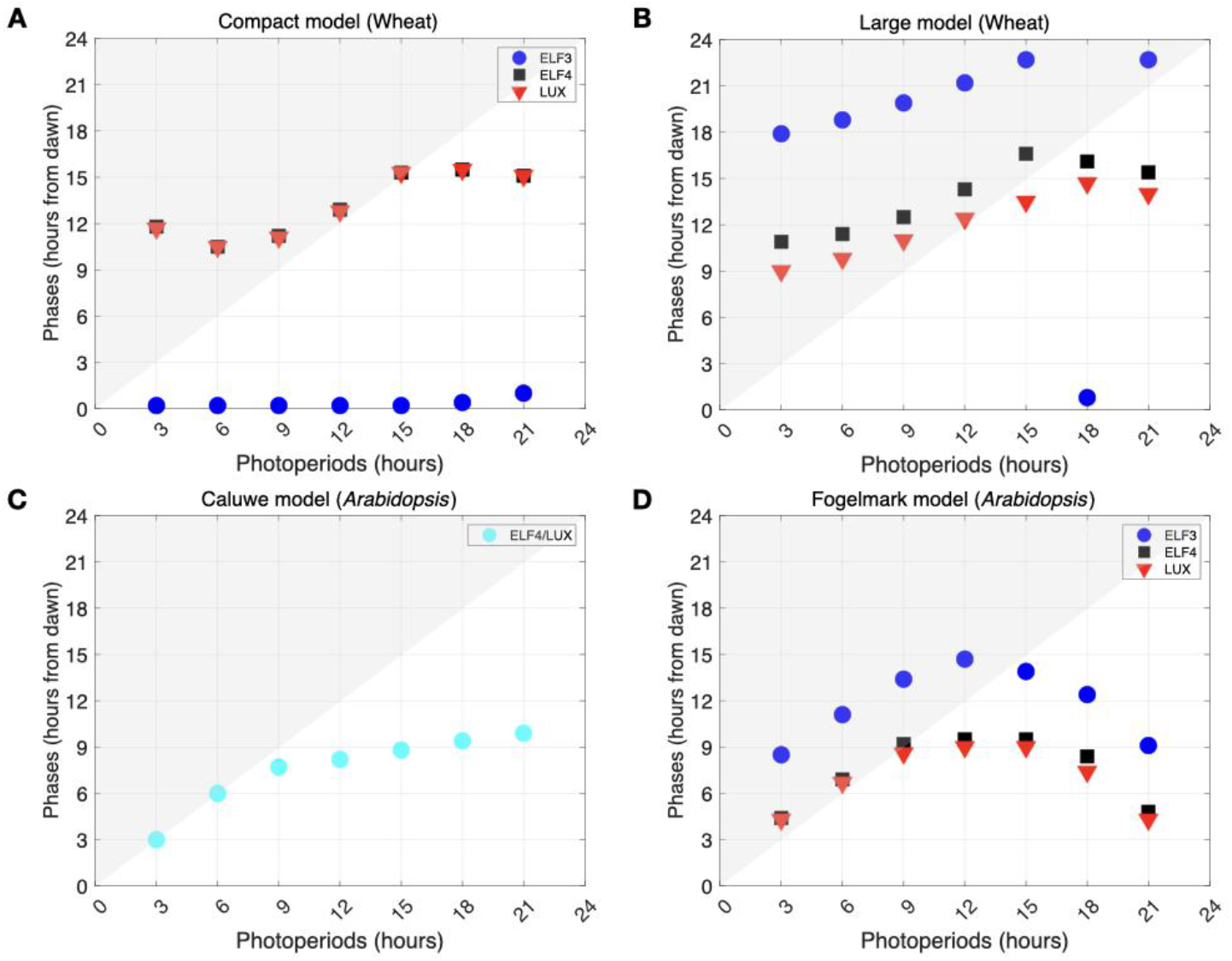
Models of wheat circadian oscillators predict *ELF3* peaks at dawn for a range of photoperiods. Phase dependence of model simulations on the photoperiod for: (A) compact wheat model, (B) large wheat model, (C) compact de Caluwe Arabidopsis model, and (D) large Fogelmark Arabidopsis model. Peak phases of *ELF3* (filled blue circle), *ELF4/LUX* (filled cyan circle), *ELF4* (filled black square) and *LUX* (filled downward red triangle) as a function of the photoperiod are simulated. A pronounced separation of dawn and dusk phases of *TtELF3* from *TtELF4* and *TtLUX* in the wheat clock models occurs for all photoperiods.

To test the robustness of the wheat oscillator models, we varied all the parameters in both models one at a time to quantify the sensitivity on period and amplitude (Fig S8). It is expected that such a parameter variation can change the circadian properties. We changed parameters by increasing and decreasing its values by 5% and evaluated the period and amplitude of *LHY* mRNA oscillations. For the large model, the parameter changes retain self-sustained oscillations, while changes lead to smoothly varying amplitudes and periods (see Fig S8). The model robustness compared favourably to that of the original large Arabidopsis network model (Fig S8, B and D). However, interestingly, for the compact wheat model, the period and oscillations were less robust than for the compact Arabidopsis model, suggesting the optimal parameter region is small (Fig S8, A and C). To further examine the robustness of both models, bifurcation analysis was performed for the parameter that determines the level of TOC1 repression on *ELF3*. In each diagram, the parameter value was varied from 0 to double of its optimised value. Both diagrams showed clear transitions from the steady states to limit cycles via Hopf bifurcations (Fig S9).

### 2.6 The wheat models predict *elf3* loss-of-function mutant dynamics

As a final test of the mathematical models of wheat circadian oscillators, we simulated the dynamics of gene expression in *elf3* loss-of-function lines. The simulated data were compared with those from *Ttelf3-A1, Ttelf3-B1* double mutant TILLING lines in the tetraploid *T. turgidum* cv Kronos lines (Wittern *et al*. 2023). As these data were not included in our parameter optimization, the simulations test the predictive power of the model.

We simulated an *elf3* mutant by reducing the translation rate of the ELF3 protein to 0.005 of its WT value, following the approach used in the original large Arabidopsis network model, where *elf3* mutants were simulated by allowing residual ELF3 expression. The large model correctly predicted low amplitude oscillations for all oscillator components in the *elf3* mutant under constant light conditions, closely mirroring experimental observations (Fig 6). In the compact wheat model, lower amplitude oscillations were correctly predicted for *LHY, LUX* and *GI*, although *TOC1* and *PRR73* oscillations were incorrectly predicted to increase in amplitude. If we abolished translation of ELF3, the large model incorrectly predicted arrhythmic clock levels under constant light conditions, although the compact model still showed a limit cycle oscillation (Fig S10). Our findings suggest that the large model can match experimental data for the *elf3* mutant if residual expression of the ELF3 protein is permitted, as was the case for the original large Arabidopsis model that it was based on.

**Figure 6.**
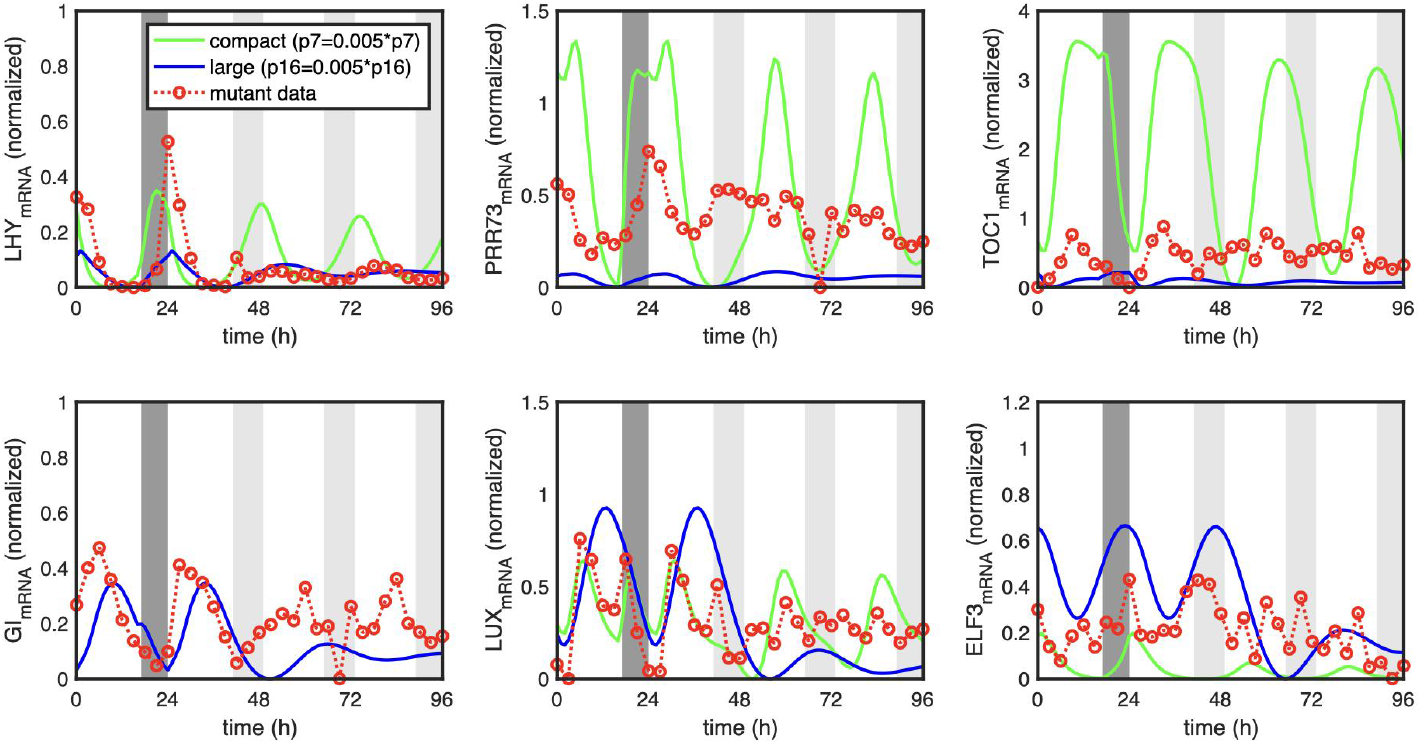
Wheat model simulations in the *elf3* mutant background. Key circadian oscillator gene dynamics in *elf3* mutant background for both compact (green solid line) and large (blue solid line) wheat models, and *Ttelf3* mutant experimental data (red dotted line). *Ttelf3* mutant is simulated by reducing the ELF3 protein translation rate to 0.005 times lower than WT. Data is normalised to wild type levels.

## 3. Discussions and conclusions

Our work presents a quantitative framework for understanding the wheat circadian oscillator. By building on fundamental insights from the model plant Arabidopsis, we developed models that capture key differences between the circadian systems of wheat and Arabidopsis.

Experimental data revealed a striking difference in the timing of *ELF3* transcript abundance between Arabidopsis and cereals (Fig 1). To explore the mechanisms underlying this divergence and its potential effects on the circadian system, we constructed two mathematical models (Fig. 2) that reproduce essential features of wild-type rhythms. These models successfully recapitulate the dawn-peaking expression dynamics of *TtELF3* observed in wheat (Figs. 3, 4). Importantly, the models were optimized using rigorous, data-driven approaches.

Our models incorporate repression of *TtELF3* transcription by TOC1, supported by the presence of TOC1 binding sites in the *TtELF3* promoter (Fig 2). Comparative analysis of the *ELF3* promoter sequences between Arabidopsis and wheat suggests that an increased number of TOC1 binding sites in wheat may underlie the observed shift in *ELF3* transcript accumulation to dawn. Our modelling results further support this as a plausible explanation, though experimental data will be required to test this hypothesis further.

Moreover, the wheat circadian oscillator models predicted that wheat *ELF3* mRNA dynamics can peak near morning for a range of photoperiods. A comparison of wheat and Arabidopsis models illustrates an uncoupling of the peak phase of *ELF3* from those of *ELF4* and *LUX* (Fig 5). Lastly, our wheat model simulations largely reproduce the experimental dynamics of *Ttelf3* mutant phenotypes (Fig 6). Further iteration between experiment and simulations will be required in future to examine other differences between the wheat and Arabidopsis circadian oscillators, but it is positive that the model was able to capture experimental details, such as the *Ttelf3* mutant phenotype and ELF3 protein dynamics, that were not included in the model optimisation.

Plant circadian oscillators as described in Arabidopsis and other species share the common structure of *CCA1/LHY* expression near dawn, *PRRs* expressed through the day to *PRR1/TOC1* at dusk and the co-expression of EC proteins at dusk. This basic structure persists in many species, though gene duplications mean that in most species the dual function of *CCA1* and *LHY* is carried out by *LHY*, and the *PRRs* are often represented by fewer members of the gene family. Plant circadian oscillators have a degree of plasticity with respect to the interval between the sequential expression of the components (CCA1/LHY-PRRs-EC) and also plasticity between the phase of expression and the phase of the external light-dark cycle (Webb *et al*. 2019). However, the shift of *ELF3* from a dusk peak in LD cycles in Arabidopsis to a dawn peak, and loss of synchrony of expression with *LUX* in wheat, is a greater level of plasticity in circadian oscillator function than was to be expected from studies on non-cereal plants.

Our simulations found that the phase of the other oscillator components can remain the same even with the dawn-phasing of *ELF3* (Figure 4). Evidence from an experimentally induced switch of mice from nocturnal to diurnal behaviour induced by altered experimental limited feeding, suggests that the timing of oscillator gene expression is rather fixed and not associated with altered outputs of behaviour, at least in the suprachiasmatic nucleus which is the timing centre of the brain. In some other tissues the diurnal behaviours were associated with earlier phasing of core oscillator genes (Challet 2007); (van Rosmalen *et al*. 2024). These data suggest that changes in the timing of oscillator gene expression are not needed for altered functions. Instead, it is the timing of output gene expression that is dramatically altered in phase to bring about altered phases of activity (van Rosmalen *et al*. 2024).

If the circadian system in wheat is analogous to those in mice, then it is possible that the change of timing of *ELF3* expression might be more associated with its role as an output of the circadian oscillator than its role in the core oscillator. *ELF3* regulates the photoperiodic regulation of flowering time (heading) in the cereals (Wittern *et al*. 2023); (Andrade *et al*. 2022); (Li *et al*. 2024). Possibly the timing of expression is critical to the regulation of flowering time, and this might be associated with degradation of ELF3 protein in the light during the day. When we examined the effect of loss of one of the A, B or D homeologue of *ELF3* we demonstrated that the circadian oscillator appears to be robust to the dosage of ELF3, whereas flowering is sensitive (Wittern *et al*. 2023). Thus, the altered timing of ELF3 may have minimal impact on oscillator function, as even low levels of ELF3 are sufficient to maintain circadian oscillations in wheat. Overall, our new models of the regulation of ELF3 abundance within wheat circadian oscillators will allow further examination of the role of ELF3 in the chronobiology of cereals, including the regulation of flowering.

## 4. Methods

### 4.1 ELF3 promoter region analysis

Promoter motifs were selected for analysis based on demonstrated binding of the DNA sequence to TOC1 or CCA1 (and/or LHY) in *A. thaliana* and/or wheat. Reference genomes analysed were *A. thaliana* genome TAIR10 (Cheng *et al*. 2017; Reiser *et al*. 2024; Swarbreck *et al*. 2008) and *T. aestivum* genome IWGSC RefSeq v1.0 (International Wheat Genome Sequencing Consortium (IWGSC) 2018). To assess the conservation of ARGF residues, which have been determined essential for the ability of AtTOC1 to bind the T1ME motif (Gendron *et al*. 2012), amino acid sequences of AtTOC1 (AT5G61380) and TaTOC1 (TraesCS6A02G227900, TraesCS6B02G253900, TraesCS6D02G207100) transcripts were aligned via multiple sequence alignment (MSA) using Clustal Omega (1.2.4) with default parameters (Madeira et al., 2024 Nucleic Acids Research).

### 4.2 Models

The Compact Wheat model was derived from the minimal models of Arabidopsis (De Caluwé *et al*. 2016; Greenwood *et al*. 2022). The large wheat model was based on a comprehensive gene-regulatory network model of Arabidopsis (Fogelmark & Troein 2014). We revised the network structure of these models by incorporating TOC1-mediated repression on *ELF3* (red lines in Fig 2A and Fig 2B). Next we explain both models in detail.

#### Compact Wheat model

We based our model on the updated version of the Arabidopsis compact model (Greenwood *et al*. 2022), which was modified from (De Caluwé *et al*. 2016) (Fig 2A). Our revised model consists of 16 ordinary differential equations (ODEs) and 3 algebraic equations (see Supplementary Section 1). The equations describe the dynamics of the mRNA and protein levels of key clock genes, represented as variables: *LHY* (LC), *PRR95, PRR73, P5T1* (*PRR59*, and *TOC1*), *ELF3, ELF4, LUX* and EC (complex of ELF3, ELF4, and LUX). The dark-accumulating protein *P*, as defined in (Locke *et al*. 2005), induces the transcription of *PRR95* and *LHY* during the transition from dark to light. Since *PRR* orthologues in wheat can not be confidently mapped directly to single genes in Arabidopsis due to independent divergence in the cereals, they were described by names that reflect the most similar genes in Arabidopsis (Murakami *et al*. 2007). This prompted us to model PRR95 and PRR73 separately, rather than combining AtPRR7 and AtPRR9 as a single species P97 in the original compact model. Moreover, we not only model ELF3 but also ELF4 and LUX as separate proteins rather than as a combined EC. We also included TOC1 repression on *ELF3*. These together are the new additions we made to the compact model of Arabidopsis circadian oscillator. The revised model contains 66 parameters (Table S1).

#### Large wheat model

The schematic shown in light blue in Fig. 2B represents the large model of the wheat circadian oscillator. This model has been extended from the work by (Fogelmark & Troein 2014) with the addition of TOC1 repression on *ELF3*. It consists of 35 ordinary differential equations (ODEs) and 10 algebraic equations (see Supplementary Section 2). The large model includes six additional oscillator genes — *REVILLE8* (*RVE8*), *GI, ZTL, CCA1, NOX/BROTHER OF LUX ARRHYTHMO* (*NOX*), and *CONSTITUTIVE PHOTOMORPHOGENIC 1* (*COP1*) — not present in the compact model. In total, it contains 123 parameters (see Table S2).

### 4.3 Parameter optimization

For both compact and large models, the parameter values were optimised by the Multi-Objective Genetic Algorithm implemented in MATLAB. One objective function measured the gaps in the peak and trough timings between the experimentally-measured (Wittern *et al*. 2023) and simulated gene expression levels. Another objective function constrained the model to oscillate with a period of 24 hr in constant light conditions and not to damp quickly under constant conditions. Among the model parameters, those related to *ELF3* dynamics were optimised.

In the compact wheat model, where possible the parameter values were initially taken from one of the top optimal parameter sets of the Greenwood revision of the De Caluwé model (De Caluwé *et al*. 2016; Greenwood *et al*. 2022). 10 parameters associated with TOC1 repression on *ELF3* and other *ELF3*-related reactions were selected and optimised.

In the large wheat model, the parameter values were taken from the values of the parameter set 2 of (Fogelmark & Troein 2014) and used as the pre-optimised values. Then, 9 parameters, e.g., TOC1 repression on *ELF3*, were optimised.

Note that, in both models, parameters, denoted by symbol w, were introduced to scale those optimised in the previous studies (De Caluwé *et al*. 2016); (Greenwood *et al*. 2022); Fogelmark & Troein 2014). They were introduced for the purpose of parameter optimisation and did not represent any biochemical constants. Details of the objective functions as well as the optimised parameter values are provided in supplementary section 3.

## Acknowledgments

This research was funded by the grant reference BB/W001209/1 of Biotechnology and Biological Sciences Research Council (BBSRC), UK. J.S.-W. was supported by the BBSRC DTP (BB/X010899/1). The authors are thankful to Will Davis and Mark Greenwood, and members of the Locke and Webb group for their fruitful discussions.

## Author contribution

Conceptualization: AW,JL; Methodology: AU,JC,AW,JL; Software: AU,JC,IT; Investigation: AU,IT,JL; Writing – Original Draft: AU; Writing – Review and Editing: AU,AW,JL; Visualisation: AU,GC,IT,AW,JL; Funding Acquisition: AW,JL.

