## Supplementary Information for "Data-driven mathematical modelling explains altered timing of *EARLY FLOWERING 3* in the wheat circadian oscillator"

**Supplemental Information**

***1: Compact Wheat Model***

The compact wheat model was based on the model developed in (De Caluwé *et al.* 2016) and further refined in (Greenwood *et al.* 2022). Based on new experimental findings, the Greenwood model of the Arabidopsis oscillator modified the De Caluwe model by replacing LHY/CCA1 activation of *PRR9/PRR7* with a repression term, by adding an autoinhibition term on *LHY/CCA1,* and also adjusted the degradation rate of PRR5/TOC1 to be higher in the dark. We thus derived our compact model of the wheat clock from the Greenwood model. In the Greenwood model, *ELF4* and *LUX* are represented as a single evening complex (*EC*) which does not explicitly include *ELF3*. To study *ELF3* behaviour, we added a new variable *ELF3*, and modelled three proteins ELF3, ELF4, and LUX separately in the EC. Due to light activation on *PRR95* but not on *PRR73*, we also modelled *PRR95* and *PRR73* separately. Parameters for *PRR95* were as for PRR97 in the Greenwood model, whilst parameters for *PRR73* were hand selected. It should also be noted that TOC1 repression on *ELF3* has not been described in models of the Arabidopsis clock. Based on *TtELF3* promoter analysis, we included TOC1 repression on *ELF3* in the compact wheat model.

The model equations are written in the form of algebraic (1-3) and ordinary differential equations (4-19) and follow a similar format to other circadian models (Burt *et al.* 2021; Pett *et al.* 2016; Singh *et al.* 2022; Upadhyay *et al.* 2019, 2020). The levels of mRNA and protein of clock component X are denoted $[X]_m$ and $[X]_p$. The rate constants for synthesis are denoted with *p*, and degradation rates for mRNA are written with *k* and protein with *d*. The protein synthesis, protein degradation and mRNA degradation terms are assumed to be linear and take the form of mass-action kinetics. The mRNA transcriptions which include activation or inhibition terms are described in the form of Hill functions (Fukuda *et al.* 2013; Locke *et al.* 2005, 2006; Partch *et al.* 2014; Pokhilko *et al.* 2013; Tokuda *et al.* 2019; Zeilinger *et al.* 2006). As in the Greenwood model, the mRNA or protein degradation rates for noon elements (PRRs, TOC1) are higher in dark than in light. Moreover, ELF4 and LUX of evening complex proteins and LHY have a higher degradation rate in light, and ELF3 degrades with the same rate in dark and light. Additionally, light is denoted by the parameter *L* (*L* = 1 during daytime; *L* = 0 during night time). Therefore, external light/dark cycles were simulated as square waves. To explore key parameter values in relation to those optimised in the preceding studies, we added three scaling parameters $w_1$, $w_2$, and $w_3$, which adjust the levels of LC, ELF3, and EC. The equations read as follows:

$$LC = w_1 * [LHY]_p \quad (1)$$

$$ELF3_{act} = w_2 * [ELF3]_p \quad (2)$$

$$EC = w_3 * [EC]_p \quad (3)$$

$$\frac{d[LHY]_m}{dt} = \frac{v_1 + v_{1L} * L * [P]_p}{1 + (\frac{LC}{K_0})^2 + (\frac{[P95]_p}{K_1})^2 + (\frac{[P5T1]_p}{K_2})^2 + (\frac{[P73]_p}{K_3})^2} - (k_{1L} * L + k_{1D} * D) * [L \quad (4)$$

$$\frac{d[LHY]_p}{dt} = p_1 + p_{1L} * L * [LHY]_m - d_1 * [LHY]_p \quad (5)$$

$$\frac{d[P]_p}{dt} = 0.3 * (1 - [P]_p) * D - d_p * [P]_p * L \quad (6)$$

$$\frac{d[P95]_m}{dt} = \frac{v_{2L} * L * [P]_p + v_{2A}}{1 + (\frac{[P5T1]_p}{K_4})^2 + (\frac{EC}{K_5})^2 + (\frac{LC}{K_{5b}})^2} - k_2 * [P95]_m \quad (7)$$

$$\frac{d[P95]_p}{dt} = p_2 * [P95]_m - (d_{2D} * D + d_{2L} * L) * [P95]_p \quad (8)$$

$$\frac{d[P73]_m}{dt} = \frac{v_3}{1 + (\frac{[P5T1]_p}{K_6})^2 + (\frac{EC}{K_7})^2 + (\frac{LC}{K_8})^2} - k_3 * [P73]_m \quad (9)$$

$$\frac{d[P73]_p}{dt} = p_3 * [P73]_m - (d_{3D} * D + d_{3L} * L) * [P73]_p \quad (10)$$

$$\frac{d[P5T1]_m}{dt} = \frac{v_4}{1 + (\frac{LC}{K_9})^2 + (\frac{[P5T1]_p}{K_{10}})^2 + (\frac{EC}{K_7})^2} - k_4 * [P5T1]_m \quad (11)$$

$$\frac{d[P5T1]_p}{dt} = p_4 * [P5T1]_m - (d_{4D} * D + d_{4L} * L) * [P5T1]_p \quad (12)$$

$$\frac{d[ELF4]_m}{dt} = \frac{v_5}{1 + (\frac{LC}{K_{11}})^2 + (\frac{[P5T1]_p}{K_{12}})^2 + (\frac{EC}{K_{13}})^2} - k_5 * [ELF4]_m \quad (13)$$

$$\frac{d[ELF4]_p}{dt} = p_5 * [ELF4]_m - (d_{5D} * D + d_{5L} * L) * [ELF4]_p - p_8 * [LUX]_p * [ELF4]_p * ELF3_{act} \quad (14)$$

$$\frac{d[LUX]_m}{dt} = \frac{v_6}{1 + (\frac{LC}{K_{14}})^2 + (\frac{[P5T1]_p}{K_{15}})^2 + (\frac{EC}{K_{16}})^2} - k_6 * [LUX]_m \quad (15)$$

$$\frac{d[LUX]_p}{dt} = p_6 * [LUX]_m - (d_{6D} * D + d_{6L} * L) * [LUX]_p - p_8 * [LUX]_p * [ELF4]_p * ELF3_{act} \quad (16)$$

$$\frac{d[ELF3]_m}{dt} = \frac{v_{3A}}{1+(\frac{[P5T1]_p}{K_{17}})^2+(\frac{LC}{K_{18}})^2} - k_7 * [ELF3]_m \tag{17}$$

$$\frac{d[ELF3]_p}{dt} = p_7 * [ELF3]_m - (d_{7D} * D + d_{7L} * L) * [ELF3]_p - p_8 * [LUX]_p * [ELF4]_p * ELF3_{act} \tag{18}$$

$$\frac{d[EC]_p}{dt} = p_8 * [LUX]_p * [ELF4]_p * ELF3_{act} - d_8 * [EC]_p \tag{19}$$

***Table S1 List of all parameter values of compact wheat model***

| Parameter name | Description | Parameter value |
|---|---|---|
| $v_1$ | LHY baseline transcription | 4.58 nM.$h^{-1}$ |
| $v_{1L}$ | LHY light-induced synthesis | 3.0 nM.$h^{-1}$ |
| $v_{2A}$ | PRR95 baseline transcription | 1.27 nM.$h^{-1}$ |
| $v_{2L}$ | PRR95 light-induced synthesis | 5.0 nM.$h^{-1}$ |
| $v_3$ | PRR73 synthesis | 1.0 nM.$h^{-1}$ |
| $\mathbf{v_{3A}}$ | ELF3 synthesis | **3.0959** nM.$h^{-1}$ |
| $v_4$ | PRR59/TOC1 synthesis | 1.1 nM.$h^{-1}$ |
| $v_5$ | ELF4 synthesis | 1 nM.$h^{-1}$ |
| $v_6$ | LUX synthesis | 2 nM.$h^{-1}$ |
| $k_{1L}$ | LHY mRNA degradation (light) | 0.53 $h^{-1}$ |
| $k_{1D}$ | LHY mRNA degradation (dark) | 0.21 $h^{-1}$ |
| $k_2$ | PRR95 mRNA degradation | 0.35 $h^{-1}$ |
| $k_3$ | PRR73 mRNA degradation | 0.56 $h^{-1}$ |
| $k_4$ | PRR59/TOC1 mRNA degradation | 0.57 $h^{-1}$ |
| $k_5$ | ELF4 mRNA degradation | 0.4 $h^{-1}$ |
| $k_6$ | LUX mRNA degradation | 0.4 $h^{-1}$ |
| $\mathbf{k_7}$ | ELF3 mRNA degradation | **0.5959** $h^{-1}$ |
| $p_1$ | LHY translation | 0.76 $h^{-1}$ |
| $p_{1L}$ | LHY light-induced translation | 0.42 $h^{-1}$ |
| $p_2$ | PRR95 translation | 1.01 $h^{-1}$ |
| $p_3$ | PRR73 translation | 0.64 $h^{-1}$ |
| $p_4$ | PRR59/TOC1 translation | 0.22 $h^{-1}$ |
| $p_5$ | ELF4 translation | 0.6 $h^{-1}$ |

| | | |
|---|---|---|
| $p_6$ | LUX translation | 0.6 $h^{-1}$ |
| $p_7$ | ELF3 translation | 0.8 $h^{-1}$ |
| **$p_8$** | LUX activity | **2.3107** $h^{-1}nM^{-2}$ |
| $d_1$ | LHY degradation | 0.68 $h^{-1}$ |
| $d_{2D}$ | PRR95 degradation (dark) | 0.5 $h^{-1}$ |
| $d_{2L}$ | PRR95 degradation (light) | 0.29 $h^{-1}$ |
| $d_{3D}$ | PRR73 degradation (dark) | 0.48 $h^{-1}$ |
| $d_{3L}$ | PRR73 degradation (light) | 0.38 $h^{-1}$ |
| $d_{4D}$ | PRR59/TOC1 degradation (dark) | 1.2 $h^{-1}$ |
| $d_{4L}$ | PRR59/TOC1 degradation (light) | 0.38 $h^{-1}$ |
| $d_{5D}$ | ELF4 degradation (dark) | 0.5 $h^{-1}$ |
| $d_{5L}$ | ELF4 degradation (light) | 0.8 $h^{-1}$ |
| $d_{6D}$ | LUX degradation (dark) | 0.5 $h^{-1}$ |
| $d_{6L}$ | LUX degradation (light) | 0.8 $h^{-1}$ |
| $d_{7D}$ | ELF3 degradation (dark) | 0.7 $h^{-1}$ |
| $d_{7L}$ | ELF3 degradation (light) | 0.7 $h^{-1}$ |
| **$d_8$** | EC degradation | **0.69558** $h^{-1}$ |
| $d_p$ | Dark accumulator protein degradation | 0.6 $h^{-1}$ |
| $K_0$ | Inhibition of LHY by LC | 2.80 nM |
| $K_1$ | Inhibition of LHY by PRR95 | 0.16 nM |
| $K_2$ | Inhibition of LHY by PRR59/TOC1 | 1.18 nM |
| $K_3$ | Inhibition of LHY by PRR73 | 0.5 nM |
| $K_4$ | Inhibition of PRR95 by PRR59/TOC1 | 0.23 nM |
| $K_5$ | Inhibition of PRR95 by EC | 0.63 nM |
| $K_{5b}$ | Inhibition of PRR95 by LC | 1.73 nM |
| $K_6$ | Inhibition of PRR73 by PRR59/TOC1 | 0.46 nM |
| $K_7$ | Inhibition of PRR73 by EC | 0.5 nM |

| | | |
|---|---|---|
| $K_8$ | Inhibition of PRR73 by LC | 0.6 nM |
| $K_9$ | Inhibition of PRR59/TOC1 by LC | 0.3 nM |
| $K_{10}$ | Inhibition of PRR59/TOC1 by PRR59/TOC1 | 1.9 nM |
| $K_{11}$ | Inhibition of ELF4 by LC | 0.3 nM |
| $K_{12}$ | Inhibition of ELF4 by PRR59/TOC1 | 0.5 nM |
| $K_{13}$ | Inhibition of ELF4 by EC | 2.0 nM |
| $K_{14}$ | Inhibition of LUX by LC | 0.3 nM |
| $K_{15}$ | Inhibition of LUX by PRR59/TOC1 | 0.5 nM |
| $\mathbf{K_{16}}$ | Inhibition of LUX by EC | **1.9810** nM |
| $\mathbf{K_{17}}$ | Inhibition of ELF3 by PRR59/TOC1 | **0.0649** nM |
| $\mathbf{K_{18}}$ | Inhibition of ELF3 by LC | **6.4184** nM |
| $\mathbf{w_1}$ | LC activity | **2.8719** |
| $\mathbf{w_2}$ | ELF3 activity | **3.1417** |
| $\mathbf{w_3}$ | EC activity | **2.7495** |

**Table S1.** Final parameters of compact model of wheat clock. The units for concentrations are tentatively assigned as nanomolar. However, simulated mRNA expressions and protein abundances are always normalized and shown in relative values. Moreover, $w_1$, $w_2$, and $w_3$ are scaling parameters. The final optimised parameters are depicted in bold.

***2: Large Wheat Model***

The large wheat model was constructed based on the one developed for Arabidopsis in Fogelmark and Troein, 2014. Please refer to Fogelmark & Troein, 2014 for a full derivation of this model, although we describe it briefly here. In contrast to the compact model, PRR59 and TOC1, as well as LHY and CCA1, were modelled separately. As shown in Fig 2B and described in main text methods, the large model also includes additional oscillator genes, *RVE8*, *GI*, *ZTL*, *NOX* and *COP1*, not present in the compact model. The formation of the Evening Complex (EC) starts with the homodimerization of ELF4. This homodimer then binds ELF3. Following the approach of the original model, we treated delays, such as those arising from the time it takes LUX to interact with ELF3, as effects that could be distributed across other processes within the model. Thus EC activity is modelled directly as a function of the levels of ELF3, ELF3-ELF4, LUX, and NOX.

The model equations are written in the form of algebraic (1-10) and ordinary differential equations (11-45). The dimensionless concentration levels of mRNA and protein of clock component X are denoted as $X_m$ and $X_p$, respectively, where *X* is an abbreviated component name explained as follows. LC: LHY-CCA1 complex; EC: Evening Complex of ELF3-ELF4-LUX-NOX; P: dark accumulator protein; P5: PRR59; P7: PRR73; P9: PRR95; E3 ELF3; E4: ELF4; E4D: ELF4 dimer; E34: ELF3-ELF4 complex; ZG: ZTL-GI complex. Non-subscript *L* and *D* denote light and darkness, respectively (*L* = 0 or *D* = 0 means off; *L* = 1 or *D* = 1 means on). When localization of a protein *X* is included in the model, cytosolic/nuclear translocation is denoted as, $X_{trans}$ . Moreover, the total proteins of complexes is denoted as $X_{tot}$ , protein production as $X_{prod}$ , protein degradation as $X_{deg}$, nuclear protein as $X_{pn}$ , and cytosolic protein as $X_{pc}$ . Following the notations of Arabidopsis large models (Fogelmark & Troein 2014; Pokhilko *et al.* 2012), we used the notation of $COP1_n$ and $COP1_d$ to represent nuclear COP1 protein in its night and day forms. For *ELF4*, *D* indicates a dimer. *LC* represents a weighted sum of CCA1 and LHY concentrations. $LC_c$ is the common term in the regulation of CCA1 and LHY transcription.

To maintain consistency with previous models (Fogelmark & Troein, 2014; Pokhilko et al., 2012), we adopted a parameter naming scheme in which each symbol reflects a biological function. Transcriptional activation and repression are governed by parameters *a* and *r* respectively. The symbol *m* is used for degradation of transcripts and proteins, *q* represents light-induced transcription, while *t* is used for protein transport, and *n* for COP1 production specifically. Additional parameters, denoted with symbol *p*, encompass a range of processes including protein synthesis, transport, degradation, and complex formation (Pokhilko *et al.* 2012, Fogelmark & Troein, 2014). Relative contributions from components within the LC and EC complexes are modulated by weighting factors f.

As in the compact model, the dark-accumulating protein *P* induces the transcription of *LHY/CCA1* and *PRR95*. A light mediated degradation is considered in LHY, CCA1, PRR73, GI, ZTL and COP1. The only change in network structure from (Fogelmark & Troein, 2014) is that we add a term representing TOC1 repression of *ELF3*. Additionally, we included three scaling parameters $w_1$, $w_2$, and $w_3$, which adjust the activity terms of LC and EC. The full model equations read as follows:

$$LC = w_1 * (LHY_p + f_5 * CCA1_p) \quad (1)$$

$$LC_c = \frac{q_1 * L * P_p + 1}{1 + (r_1 * P9_p)^2 + (r_2 * P7_p)^2 + (r_3 * P5_{pn})^2 + (r_4 * TOC1_{pn})^2} \quad (2)$$

$$EC = \frac{w_2 * (LUX_p + f_6 * NOX_p) * w_3 * (E34 + f_1 * E3_p)}{1 + f_3 * (LUX_p + f_2 * NOX_p) + f_4 * w3 * (E34 + f_1 * E3_p)} \quad (3)$$

$$P5_{trans} = t_5 * P5_{pc} - t_6 * P5_{pn} \quad (4)$$

$$TOC1_{trans} = t_7 * TOC1_{pc} - \frac{t_8}{1 + m_{37} * P5_{pn}} * TOC1_{pn} \quad (5)$$

$$E34_{prod} = p_{25} * E3_p * E4D \quad (6)$$

$$E3_{deg} = m_{30} * COP1_d + m_{29} * COP1_n + m_9 + m_{10} * GI_{pn} \quad (7)$$

$$ZG_{prod} = p_{12} * ZTL_p * GI_{pc} - (p_{13} * D + p_{10} * L) * ZG_p \quad (8)$$

$$E3_{tot} = E3_p + E34 \quad (9)$$

$$GI_{trans} = p_{28} * GI_{pc} - \frac{p_{29}}{1 + t_9 * E3_{tot}} * GI_{pn} \quad (10)$$

$$\frac{dLHY_m}{dt} = \frac{LC_c}{1 + (r_{11} * LC)^2} - m_1 * LHY_m \quad (11)$$

$$\frac{dLHY_p}{dt} = (L + m_4 * D) * LHY_m - m_3 * LHY_p \quad (12)$$

$$\frac{dCCA1_m}{dt} = LC_c - m_1 * CCA1_m \quad (13)$$

$$\frac{dCCA1_p}{dt} = (L + m_4 * D) * CCA1_m - m_3 * CCA1_p \quad (14)$$

$$\frac{dP_p}{dt} = p_7 * D * (1 - P_p) - m_{11} * L * P_p \quad (15)$$

$$\frac{dP9_p}{dt} = P9_m - m_{13} * P9_p \quad (16)$$

$$\frac{dP7_p}{dt} = P7_m - (m_{15} + m_{23} * D) * P7_p \quad (17)$$

$$\frac{dP5_{pc}}{dt} = P5_m - (m_{17} + m_{24} * ZTL_p) * P5_{pc} - P5_{trans} \quad (18)$$

$$\frac{dP5_{pn}}{dt} = P5_{trans} - m_{42} * P5_{pn} \quad (19)$$

$$\frac{dTOC1_{pn}}{dt} = TOC1_{trans} - \frac{m_{43}}{1 + m_{38} * P5_{pn}} * TOC1_{pn} \quad (20)$$

$$\frac{dTOC1_{pc}}{dt} = TOC1_m - (m_8 + m_6 * ZTL_p) * TOC1_{pc} - TOC1_{trans} \quad (21)$$

$$\frac{dE4_p}{dt} = p_{23} * E4_m - m_{35} * E4_p - (E4_p)^2 \quad (22)$$

$$\frac{dE4D}{dt} = (E4_p)^2 - m_{36} * E4D - E34_{prod} \quad (23)$$

$$\frac{dE3_m}{dt} = \frac{1}{(1+(r_{21}*LC)^2)*(1+(r_{41}*TOC1_{pn})^2)} - m_{26} * E3_m \quad (24)$$

$$\frac{dE3_p}{dt} = p_{16} * E3_m - E34_{prod} - E3_{deg} * E3_p \quad (25)$$

$$\frac{dE34}{dt} = E34_{prod} - m_{22} * E34 * E3_{deg} \quad (26)$$

$$\frac{dLUX_p}{dt} = LUX_m - m_{39} * LUX_p \quad (27)$$

$$\frac{dCOP1_{pc}}{dt} = n_5 - p_6 * COP1_{pc} - m_{27} * COP1_{pc} * (1 + p_{15} * L) \quad (28)$$

$$\frac{dCOP1_n}{dt} = p_6 * COP1_{pc} - (n_{14} + n_6 * L * P_p) * COP1_n - (1 + p_{15} * L) * m_{27} * COP1_n \quad (29)$$

$$\frac{dCOP1_d}{dt} = (n_{14} + n_6 * L * P_p) * COP1_n - m_{31} * (1 + m_{33} * D) * COP1_d \quad (30)$$

$$\frac{dZTL_p}{dt} = p_{14} - ZG_{prod} - m_{20} * ZTL_p \quad (31)$$

$$\frac{dZG_p}{dt} = ZG_{prod} - m_{21} * ZG_p \quad (32)$$

$$\frac{dGI_{pc}}{dt} = p_{11} * GI_m - ZG_{prod} - GI_{trans} - m_{19} * GI_{pc} \quad (33)$$

$$\frac{dGI_{pn}}{dt} = GI_{trans} - m_{19} * GI_{pn} - m_{25} * (E3_{tot}) * (1 + m_{28} * COP1_d + m_{32} * COP1_n) * GI_{pn} \quad (34)$$

$$\frac{dNOX_m}{dt} = \frac{1}{(1+(r_{28}*LC)^2)*(1+(r_{29}*P7_p)^2)} - m_{44} * NOX_m \quad (35)$$

$$\frac{dNOX_p}{dt} = NOX_m - m_{45} * NOX_p \quad (36)$$

$$\frac{dRVE8_m}{dt} = \frac{1}{(1+(r_{30}*P9_p)^2)*(r_{31}*P7_p)^2)+(r_{32}*P5_{pn})^2)} - m_{46} * RVE8_m \quad (37)$$

$$\frac{dRVE8_p}{dt} = RVE8_m - m_{47} * RVE8_p \quad (38)$$

$$\frac{dP9_m}{dt} = q_3 * L * P_p - m_{12} * P9_m + \frac{1+a_3*r_{33}*RVE8_p}{(1+r_{33}*RVE8_p)*(1+(r_5*LC)^2)*(1+(r_6*EC)^2)*(1+(r_7*TOC1_{pn})^2)*(1+(r_{40}*P5_{pn})^2)} \quad (39)$$

$$\frac{dP7_m}{dt} = \frac{1}{(1+(r_8*LC)^2)*(1+(r_9*EC)^2)*(1+(r_{10}*TOC1_{pn})^2)*(1+(r_{40}*P5_{pn})^2)} - m_{14} * P7_m \quad (40)$$

$$\frac{dP5_m}{dt} = \frac{1+(a_4*r_{34}*RVE8_p)}{(1+r_{34}*RVE8_p)*(1+(r_{12}*LC)^2)*(1+(r_{13}*EC)^2)*(1+(r_{14}*TOC1_{pn})^2)} - m_{16} * P5 \text{[illegible]} \quad (41)$$

$$\frac{dTOC1_m}{dt} = \frac{1+(a_5*r_{35}*RVE8_p)}{(1+r_{35}*RVE8_p)*(1+(r_{15}*LC)^2)*(1+(r_{16}*EC)^2)*(1+(r_{17}*TOC1_{pn})^2)} - m_5 * T \text{[illegible]} \quad (42)$$

$$\frac{dE4_m}{dt} = \frac{1+(a_6*r_{36}*RVE8_p)}{(1+r_{36}*RVE8_p)*(1+(r_{18}*EC)^2)*(1+(r_{19}*LC)^2)*(1+(r_{20}*TOC1_{pn})^2)} - m_7 * E4_m \quad (43)$$

$$\frac{dLUX_m}{dt} = \frac{1+(a_7*r_{37}*RVE8_p)}{(1+r_{37}*RVE8_p)*(1+(r_{22}*EC)^2)*(1+(r_{23}*LC)^2)*(1+(r_{24}*TOC1_{pn})^2)} - m_{34} * L \text{[illegible]} \quad (44)$$

$$\frac{dGI_m}{dt} = \frac{1+(a_8*r_{38}*RVE8_p)}{(1+r_{38}*RVE8_p)*(1+(r_{25}*EC)^2)*(1+(r_{26}*LC)^2)*(1+(r_{27}*TOC1_{pn})^2)} - m_{18} * GI_m \quad (45)$$

*Table S2 List of all parameter values of large wheat model*

| LARGE MODEL | | | | | |
|---|---|---|---|---|---|
| ***Name*** | ***Values*** | ***Name*** | ***Values*** | ***Name*** | ***Values*** |
| $a_3$ | 1.022 | $m_{31}$ | 0.3 $h^{-1}$ | $r_9$ | 0.5885 |
| $a_4$ | 9.866 | $m_{32}$ | 5.681 $h^{-1}$ | $r_{10}$ | 21.8 |
| $a_5$ | 5.353 | $m_{33}$ | 13 $h^{-1}$ | $r_{11}$ | 2.086 |
| $a_6$ | 2.278 | $m_{34}$ | 0.1115 $h^{-1}$ | $r_{12}$ | 6.033 |
| $a_7$ | 5.917 | $m_{35}$ | 0.9188 $h^{-1}$ | $r_{13}$ | 1.053 |
| $a_8$ | 4.365 | $m_{36}$ | 0.5711 $h^{-1}$ | $r_{14}$ | 12.66 |
| **$f_1$** | **0.3598** | $m_{37}$ | 0.9391 $h^{-1}$ | $r_{15}$ | 6.743 |
| $f_2$ | 2.021 | $m_{38}$ | 7.975 $h^{-1}$ | $r_{16}$ | 0.1519 |
| $f_3$ | 0.03313 | $m_{39}$ | 0.216 $h^{-1}$ | $r_{17}$ | 5.199 |
| $f_4$ | 0.1 | $m_{42}$ | 0.3759 $h^{-1}$ | $r_{18}$ | 1.205 |
| $f_5$ | 0.3853 | $m_{43}$ | 0.5214 $h^{-1}$ | $r_{19}$ | 16.24 |
| $f_6$ | 0.2525 | $m_{44}$ | 0.3577 $h^{-1}$ | $r_{20}$ | 0.1465 |
| $m_1$ | 0.996 $h^{-1}$ | $m_{45}$ | 0.7944 $h^{-1}$ | **$r_{21}$** | **0.3561** |
| $m_3$ | 0.5889 $h^{-1}$ | $m_{46}$ | 0.7541 $h^{-1}$ | **$r_{22}$** | **2.3097** |
| $m_4$ | 0.3761 $h^{-1}$ | $m_{47}$ | 0.1293 $h^{-1}$ | $r_{23}$ | 7.1 |
| $m_5$ | 2.3 $h^{-1}$ | $n_5$ | 0.23 $h^{-1}$ | $r_{24}$ | 16.33 |
| $m_6$ | 0.0133 $h^{-1}$ | $n_6$ | 20 $h^{-1}$ | $r_{25}$ | 1.027 |
| $m_7$ | 0.6487 $h^{-1}$ | $n_{14}$ | 0.1 $h^{-1}$ | $r_{26}$ | 5.466 |
| $m_8$ | 5.437 $h^{-1}$ | $p_6$ | 0.6 $h^{-1}$ | $r_{27}$ | 6.864 |
| $m_9$ | 0.1225 $h^{-1}$ | $p_7$ | 0.3 $h^{-1}$ | $r_{28}$ | 8.392 |
| $m_{10}$ | 0.01001 $h^{-1}$ | $p_{10}$ | 0.2 $h^{-1}$ | $r_{29}$ | 0.1423 |
| $m_{11}$ | 0.6771 $h^{-1}$ | $p_{11}$ | 1.78 $h^{-1}$ | $r_{30}$ | 2.714 |

| | | | | | |
|---|---|---|---|---|---|
| $m_{12}$ | $1.988\ h^{-1}$ | $p_{12}$ | $8\ h^{-1}$ | $r_{31}$ | 0.01041 |
| $m_{13}$ | $0.376\ h^{-1}$ | $p_{13}$ | $0.7\ h^{-1}$ | $r_{32}$ | 4.775 |
| $m_{14}$ | $4.916\ h^{-1}$ | $p_{14}$ | $0.3\ h^{-1}$ | $r_{33}$ | 0.9026 |
| $m_{15}$ | $0.09303\ h^{-1}$ | $p_{15}$ | $3\ h^{-1}$ | $r_{34}$ | 0.05704 |
| $m_{16}$ | $0.5828\ h^{-1}$ | $p_{16}$ | $0.4024\ h^{-1}$ | $r_{35}$ | 0.02929 |
| $m_{17}$ | $0.04744\ h^{-1}$ | $p_{23}$ | $1.461\ h^{-1}$ | $r_{36}$ | 0.49 |
| $m_{18}$ | $2.426\ h^{-1}$ | $p_{25}$ | $1.111\ h^{-1}$ | $r_{37}$ | 0.554 |
| $m_{19}$ | $0.2\ h^{-1}$ | $p_{28}$ | $2.13\ h^{-1}$ | $r_{38}$ | 0.05062 |
| $m_{20}$ | $1.8\ h^{-1}$ | $p_{29}$ | $25.2\ h^{-1}$ | $r_{40}$ | 1.051 |
| $m_{21}$ | $0.1\ h^{-1}$ | $q_{1}$ | $0.1217\ h^{-1}$ | $t_{5}$ | 1.103 |
| $m_{22}$ | $0.3012\ h^{-1}$ | $q_{3}$ | $0.2873\ h^{-1}$ | $t_{6}$ | 0.5891 |
| $m_{23}$ | $0.1764\ h^{-1}$ | $r_{1}$ | 3.747 | $t_{7}$ | 0.2317 |
| $m_{24}$ | $2.848\ h^{-1}$ | $r_{2}$ | 2.384 | $t_{8}$ | 0.1472 |
| $m_{25}$ | $0.4176\ h^{-1}$ | $r_{3}$ | 4.747 | $t_{9}$ | 0.8543 |
| **$m_{26}$** | **$0.1876\ h^{-1}$** | $r_{4}$ | 16.3 | **$w_{1}$** | **0.7723** |
| $m_{27}$ | $0.1\ h^{-1}$ | $r_{5}$ | 0.1 | **$w_{2}$** | **0.1466** |
| $m_{28}$ | $0.02757\ h^{-1}$ | $r_{6}$ | 0.4728 | **$w_{3}$** | **0.5692** |
| $m_{29}$ | $0.01\ h^{-1}$ | $r_{7}$ | 35.94 | **$r_{41}$** | **27.6647** |
| **$m_{30}$** | **$1.0731\ h^{-1}$** | $r_{8}$ | 2.225 | $v_{3A}$ | 1 |

**Table S2.** Final parameters of the large model of wheat clock. The optimised parameters are depicted in bold.

***3: Parameter Optimisation***

We optimised the parameters by fitting the model to the peaks and troughs of RNA abundance data for key wheat clock components (Wittern et al., 2023). The time series of RNA abundance data (Wittern et al., 2023) were smoothed using the "smooth" function of the curve fitting toolbox in MATLAB to remove noisy spikes in expression (see Fig S2). The peak and trough times were identified from the experimental and simulated data using the "findpeaks" function in MATLAB. The peaks at time 0 h that were not identified by “findpeaks'' were added manually (Fig S3). In the optimization, two objective functions were minimised:

**Objective Function 1 (*ObjFun1*)**

$$ObjFun1 = \sum_{i=1}^{VarTotal} \left( \sum_{j=1}^{Peaks_i} |Peaks_{ij}^{Exp} - Peaks_{ij}^{Model}| + \sum_{j=1}^{Troughs_i} |Troughs_{ij}^{Exp} - Troughs_{ij}^{Model}| \right)$$

Where:

*i* indexes each variable (Gene), with a total of *VarTotal* genes examined

*j* indexes each detected peak or trough.

$Peaks_{ij}^{Exp}$ $and$ $Troughs_{ij}^{Exp}$ are the times of the experimentally measured peaks and troughs.

$Peaks_{ij}^{Model}$ and $Troughs_{ij}^{Model}$ are the corresponding times from the model.

The objective function 1 (*ObjFun1*) was used to match the peak and trough timings between the experimentally-measured and simulated gene expressions. This was performed over all peaks and troughs to capture the period lengthening observed in constant light conditions in the experimental data. The absolute distance between the timings of experimentally-measured and simulated peaks (or troughs) with the closest phase was calculated. The distance was then averaged across peaks (or troughs) and summed across all variables. To emphasise the dawn peak of *ELF3*, more weight was put on the terms for the *ELF3* than those for the other genes. To request more precise fitting in an LD condition, more weight was put on the terms in an LD condition than those in a constant light condition. In addition, to avoid clock genes showing double peaks (or troughs), a penalty was given if more than two peaks (or troughs) were detected within a window of 7 hr. If no peak (or trough) was detected in the model, a large penalty was given.

**Objective Function 2 (*ObjFun2*)**

$$ObjFun2 = | - log(\nabla_{last12hr} + 10^{-70}) | + |Period_{LHY}^{Model} - 24|$$

The objective function 2 (*ObjFun2*) constrained the model to ensure that circadian components oscillate with a period of 24 hr in constant light and the oscillation does not damp quickly under constant conditions. It is a necessary condition for a circadian gene to have an autonomous rhythm. The first term tested the oscillatory activity at the end of the constant light simulation by penalising a very small change in the clock gene oscillation. For the compact model, $\nabla_{last12hr}$ was calculated as $\nabla_{last12hr} = | [ELF3]_m(t_{end}) - [ELF3]_m(t_{end} - 12) | /12$. The second term measured deviation of the simulated oscillation period of *LHY* from 24 hr.

**Parameter Selection and Optimisation**

Among the model parameters, those related to *ELF3* dynamics, *e.g.*, TOC1 repression on *ELF3*, were selected and optimised. In the compact model, 10 parameters ($w_1$, $w_2$, $w_3$, $v_{3A}$, $p_8$, $k_7$, $d_8$, $K_{16}$, $K_{17}$, and $K_{18}$) were selected, while 9 parameters ($w_1$, $w_2$, $w_3$, $r_{41}$, $f_1$, $m_{26}$, $m_{30}$, $r_{21}$, and $r_{22}$) were selected in the large model. In both models, the parameters, $w_1$, $w_2$, and $w_3$ were used to scale the optimised values of the previous studies. They were introduced for the purpose of parameter optimization and did not represent any biochemical constants (i.e., dimensionless).

The two objective functions were simultaneously optimised using the Multiple-Objective Genetic Algorithm from the Global Optimization Toolbox ("gamultiobj" package) in MATLAB. After multiple rounds of optimizations, final parameter values were obtained. The optimised parameter values can be found in bold letters in Tables S1 and S2.

*Figure S1 ELF3 promoter region analysis comparison in monocot species, including cereals*

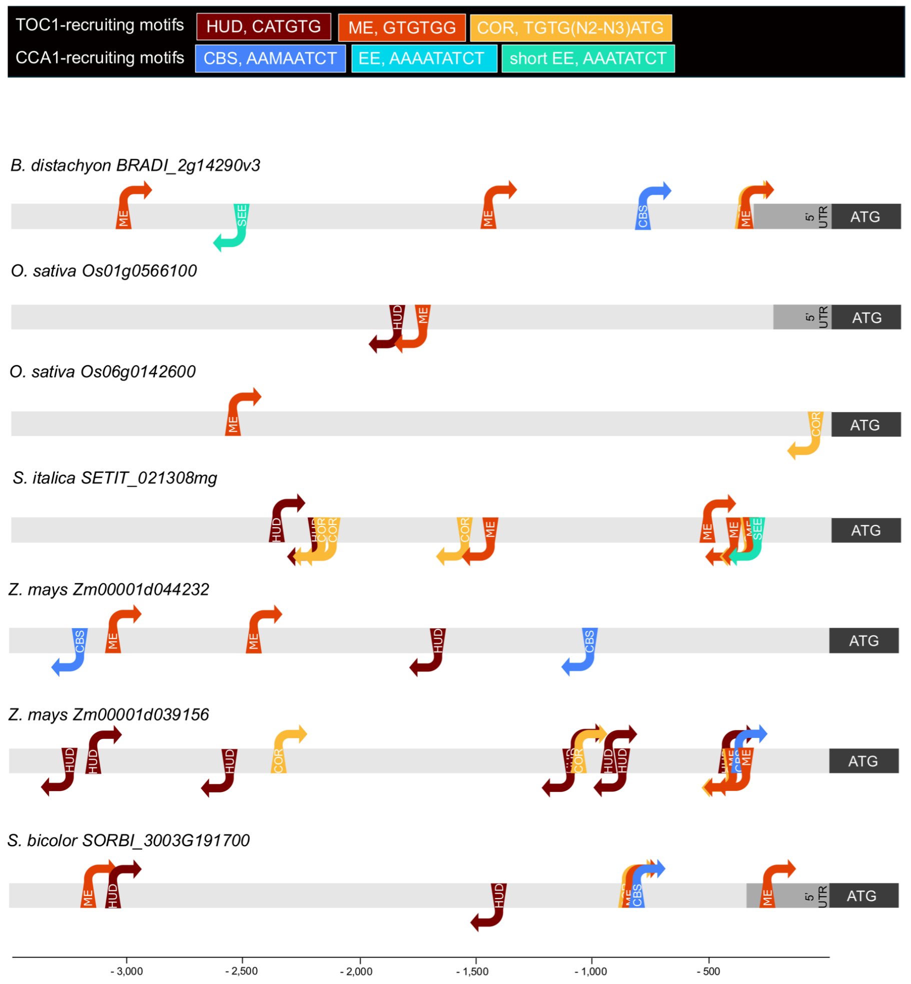


**Figure S1.** ***ELF3 promoter region analysis comparison in monocot species, including cereals.*** Representation of genomic DNA sequences 3500 base pairs upstream of the start codon of At*ELF3* orthologs in monocots, including a number of cereals . Motifs were selected for analysis based on demonstrated binding of TOC1 or CCA1/LHY. TOC1-recruiting motifs include the Morning Element (MEGTGTGG) (Harmer & Kay 2005), Hormone Up-regulated at Dawn motif (HUD, CATGTG) (Michael *et al.* 2008); (Gendron *et al.* 2012), and CO-response element (COR, TGTG(N2-N3)ATG, (Gendron *et al.* 2012); (Tiwari *et al.* 2010). CCA1/LHY-binding motifs include the Evening Element (EE, AAAATATCT) (Alabadí *et al.* 2001; Harmer & Kay 2005; Harmer *et al.* 2000), short EE element (SEE, AAATATCT) (Gong *et al.* 2022), CCA1 binding site (AAAAATCT or AACAATCT) (Wang *et al.* 1997); (Lu *et al.* 2012). Sequences were analysed from reference genomes available on Ensembl Plants (plants.ensembl.org) and the Gene ID of the *ELF3* ortholog is shown in the figure. Species and reference genomes analysed: *Oryza sativa*, genome IRGSP-1.0 (International Rice Genome Sequencing Project 2005; Kawahara *et al.* 2013); *Brachypodium distachyon*, genome Brachypodium_distachyon_v3.0 (Fox *et al.* 2013; International Brachypodium Initiative 2010); *Setaria italica*, genome Setaria_italica_v2.0 (Bennetzen *et al.* 2012; Zhang *et al.* 2012); *Sorghum bicolor*, genome Sorghum_bicolor_NCBIv3 (McCormick *et al.* 2018; Paterson *et al.* 2009); *Zea mays*, genome Zm-B73-REFERENCE-NAM-5.0 (Hufford *et al.* 2021; Jiao *et al.* 2017).

***Figure S2 Data normalisation 1: smoothing step***

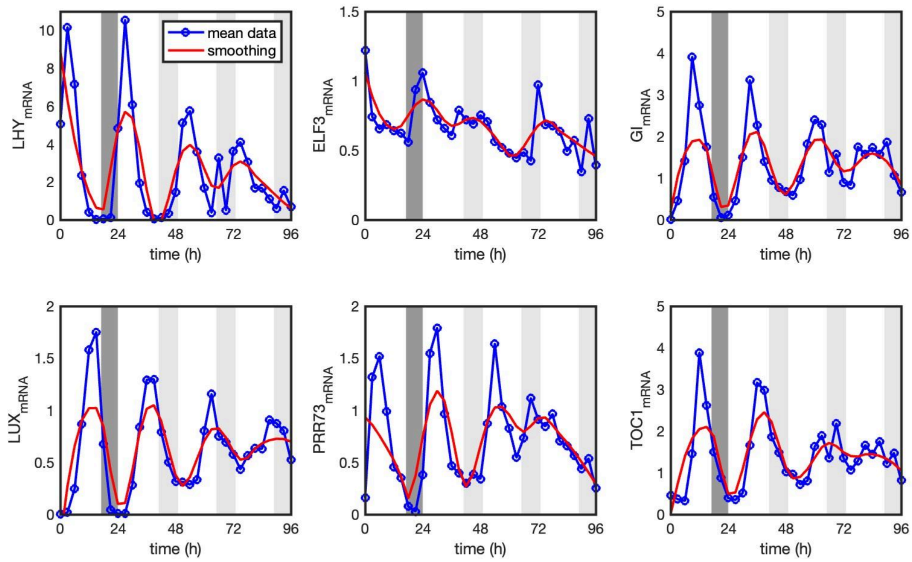


**Figure S2.** ***Data normalisation 1: smoothing step*****.** Measured changes in wheat oscillator gene RNA abundance data (blue line) was smoothed (red line) using MATLAB's inbuilt function "smooth" before optimization. This smoothed data was used for peak calculations to compare to model simulations.

### *Figure S3 Data normalisation 2: peak calling*

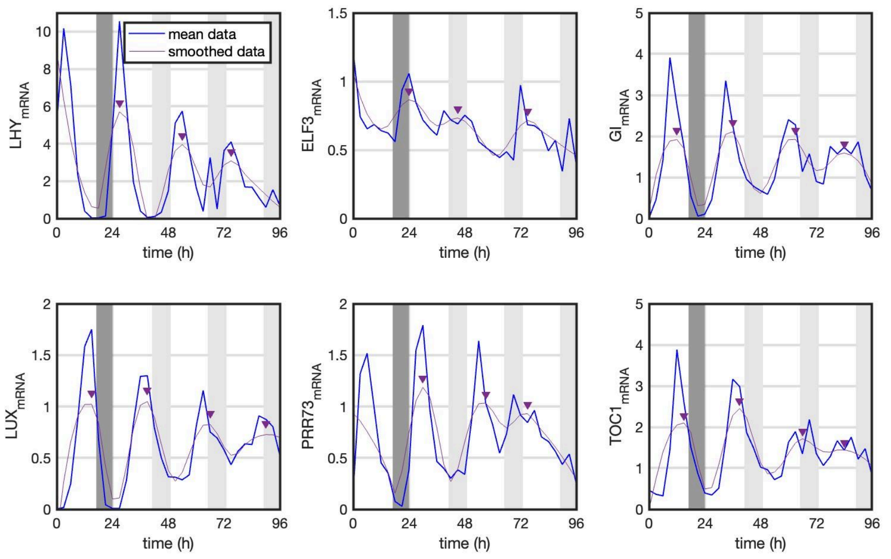


**Figure S3.** ***Data normalisation 2: peak calling.*** Peak phases for wheat oscillator gene transcripts (blue line) were calculated using MATLAB's inbuilt function "findpeaks" on smoothed data (red line). Peaks (red values) were plugged into an objective function (ObjFun1) for the optimization. In the case of *LHY* and *PRR73* where the first peak was not captured on a smoothed curve (red line) but present in raw data (blue line), we added the first peaks.

### *Figure S4* **ELF3** *protein degrades faster under light in Large model*

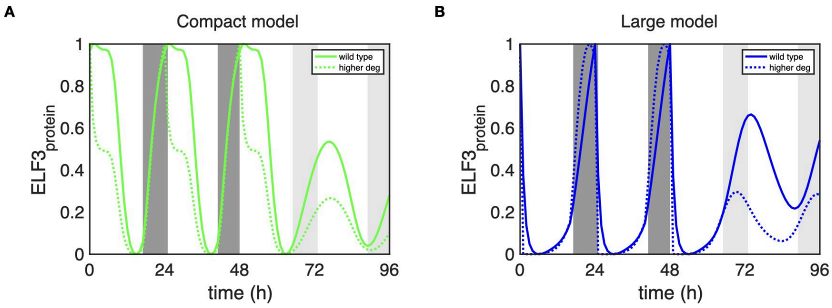


**Figure S4. ELF3** ***protein degrades faster under light in Large model.*** ELF3 protein dynamics in the compact (**A**) and large **(B)** wheat model simulations under 48 hours of light-dark cycles followed by 48 hours of constant light. A sharp drop in ELF3 protein levels are observed at the onset of light in the large model but not the compact model. The protein abundance reduces more abruptly when light-mediated degradation rates are increased (dashed lines) when compared to wild type (in solid lines). Light-mediated ELF3 degradation rates, $d_{7L}$ in compact and $m_{30}$ in large model, have been increased by 2 fold and 10 fold, respectively.

### *Figure S5* *Reoptimization can not rescue ELF3 dynamics*

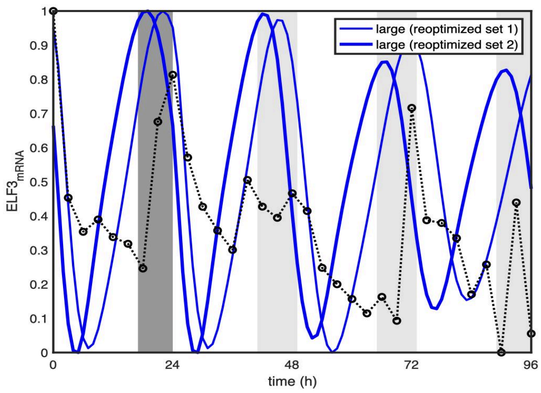


**Figure S5. Reoptimization can not rescue** ***ELF3*** **dynamics.** Re-optimisation was performed without TOC1 repression of *ELF3*. Here for example, for a large model, *ELF3* mRNA dynamics with two sets (lighter and darker blue lines) of re-optimised values (of the same 9 parameters which were used in optimisation) is shown.

### *Figure S6* *Removal of* CCA1 *repression of ELF3 does not affect simulated clock rhythms*

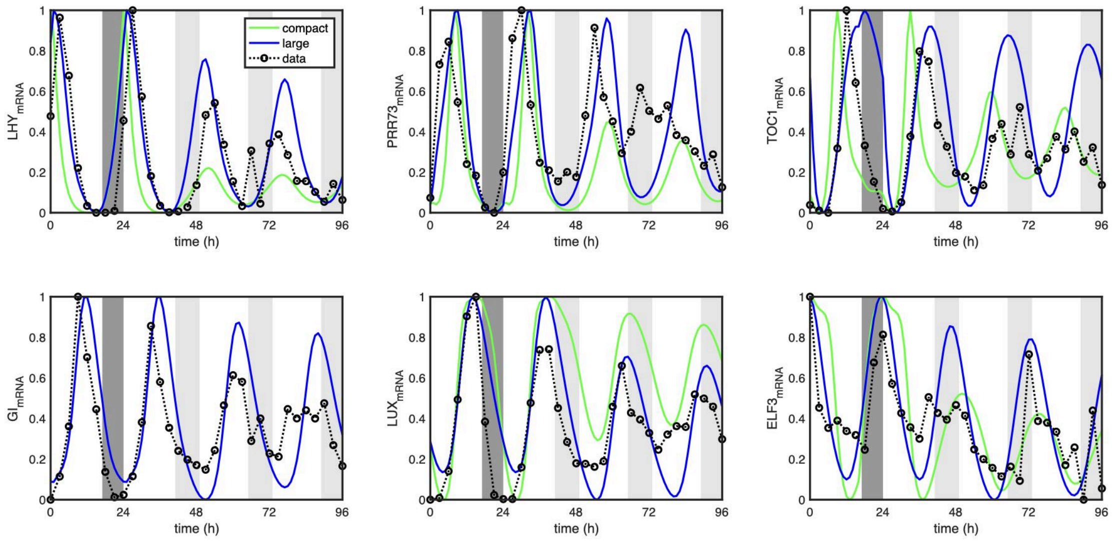


**Figure S6.** ***Removal of*** **CCA1** ***repression of ELF3 term does not affect simulated clock rhythms*** Model simulations after removal of term responsible for CCA1 repression on *ELF3* leads to almost no change in clock rhythms. All clock genes retain their dynamics except subtle waveform change on *ELF3* in the LD cycle for the compact model (in green). To minimise CCA1 repression of *ELF3* in the compact model we set $K_{18}$ 10 times higher and in the large model we set $r_{21}$ to 0.

### *Figure S7* Simulated expression of wheat clock genes under 3 different photoperiods

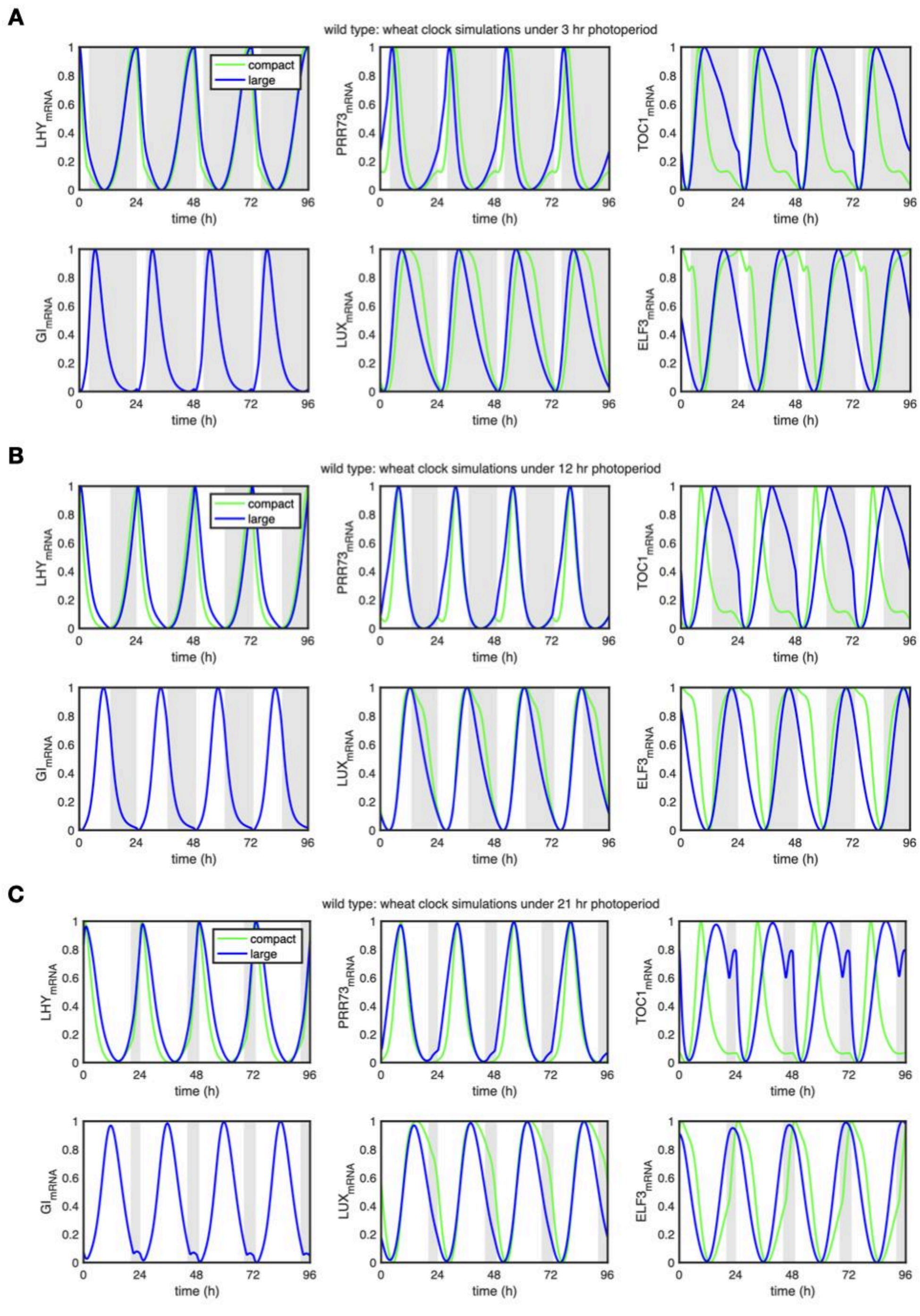


**Figure S7. Simulated expression of wheat clock genes under 3 different photoperiods**. Three photoperiods from Fig 5 are selected to visualise the gene expression dynamics. The compact model (in green) and large model (in blue) are shown under **(A)** short day photoperiod (3 hour light and 21 hour darkness), **(B)** normal day (12:12) and **(C)** longer day (21:3). White panel depicts light and the grey panel is dark.

### *Figure S8 Sensitivity analysis*

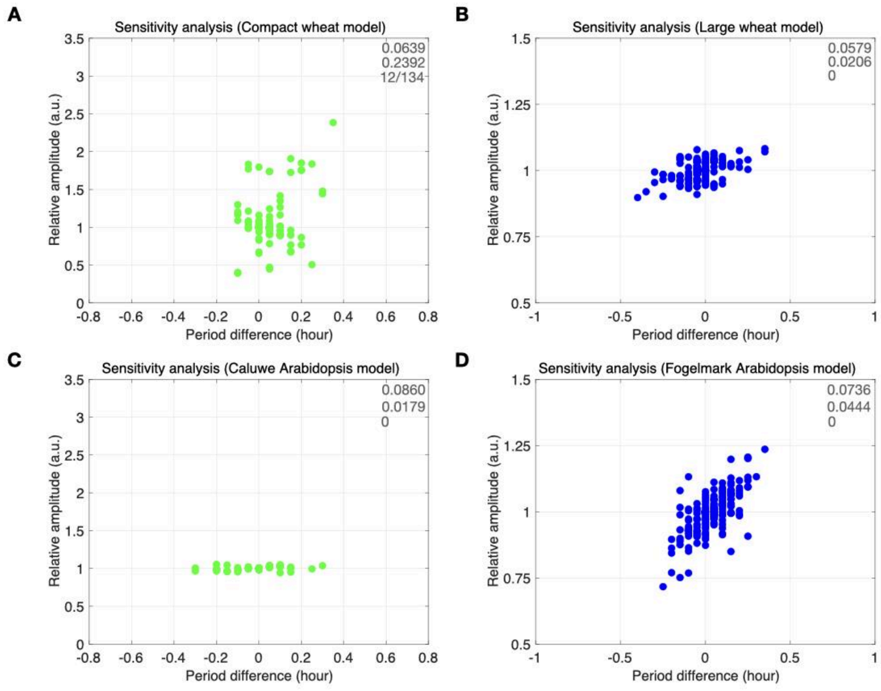


**Figure S8. Sensitivity analysis for the wheat and Arabidopsis clock models.** Sensitivity analysis was carried out by quantifying amplitude and period changes upon varying each parameter by 5%, one at a time, for the wheat compact (A), wheat large (B), Arabidopsis Caluwe (C), Arabidopsis Fogelmark (D) models, respectively. Each parameter was increased and decreased by 5% of its value and corresponding amplitude and period of the system was calculated using the *LHY* mRNA variable. Periods differences (period after perturbation - default period) are denoted on x-axis and relative amplitude (amplitude after perturbation/default amplitude) on y-axis with each parameter as circular dots (green and blue for compact and large models, in A, C and B, D panel respectively). The three numbers on top-right of each subfigure show the mean of periods and amplitudes, and number of parameters for which undefined values were obtained, respectively.

*__Figure S9 Bifurcation analysis__*

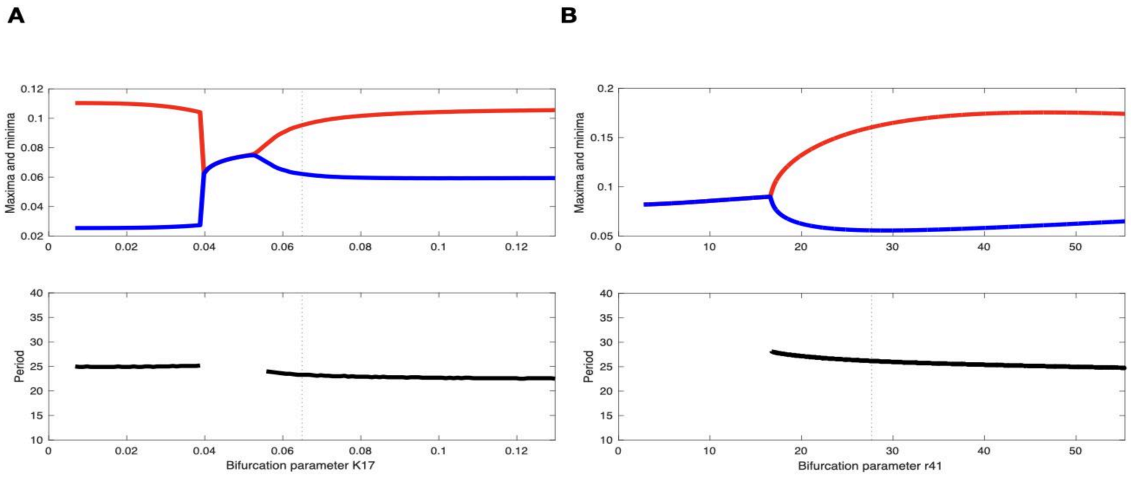


**Figure S9.** ***Bifurcation analysis.*** Bifurcation diagrams were generated for amplitude and period of one of the optimised free parameters, TOC1 repression on *ELF3*, of both wheat models, $K_{17}$ of compact **(A)** and $r_{41}$ of large **(B)**, respectively. Bifurcation parameter is varied from 0 to double of its default value (denoted at vertical dashed line) on the x-axis. On top panels (in A & B), maxima (in red curved line) and minima (in blue curved line) of limit cycle oscillations were calculated using the variable *LHY*. Steady state with no oscillation zone (single blue line) is shown where maxima and minima collide. Moreover, the bottom panel (in A & B) shows the period of the system (in black line) in the oscillatory regime.

***Figure S10 Dynamics of model simulations under the complete elf3 knockout***

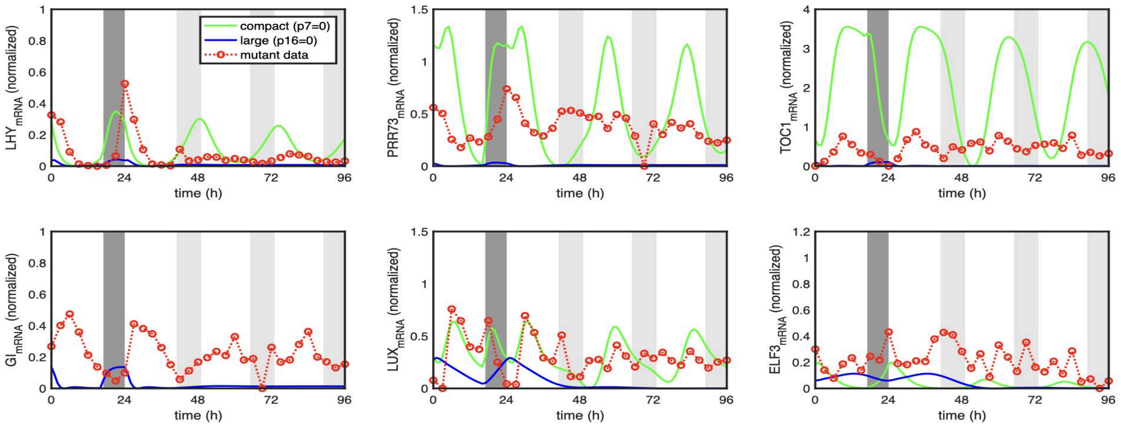


**Figure S10.** ***Dynamics of model simulations under the complete elf3 knockout.*** The *elf3* mutant dynamics simulated in the compact and large wheat models (green and blue lines) by setting the ELF3 protein translation rates $p_7$ and $p_{16}$ to 0, respectively. Experimental *elf3* data is shown in a red dotted line with empty circles, the same as in Fig 6.
